# Autophagy driven by VPS34 enables differentiated cell plasticity and cancer initiation

**DOI:** 10.1101/2025.05.02.651270

**Authors:** F. Ramos-Delgado, H. Shalhoub, C. Guyon, C. Handschin, P. Cerapio-Arroyo, R. D’Angelo, N. Therville, A. Villard, C. Cayron, C. Valle, E. Sarot, N Dussere, M. Di-Luoffo, V. Rebours, A. Couvelard, C. Joffre, H. de Oliveira, M. Dufresne, B. Thibault, J. Guillermet-Guibert

## Abstract

Differentiated cell plasticity and autophagy are fundamental mechanisms of tissue repair. To investigate their integration to maintain tissue homeostasis, we developed a unique model in which VPS34, the highly conserved class III PI3K that promotes autophagic flux, was inactivated in pancreatic exocrine cells, recognised study models. Using scRNAseq, we found that VPS34-null acinar cells evolved towards a transcriptional identity present at low levels in mouse and human pancreas. Functionally, it unexpectedly prevented transdifferentiation, chronic pancreatitis and precancer initiation triggered by stressors. Mechanistically, the newly differentiated cell state was less susceptible to cancer promotion by oncogenic KRAS, through reduced class I PI3K and increased lysosomal degradation of pro-inflammatory REG3A. This result opens the so far unreachable possibility of strategies for protection from environmentally induced cancer initiation.

## Main Text

Cell plasticity is a fundamental mechanism of tissue homeostasis, particularly in tissues that lack stem cells. This protective mechanism of environmentally induced tissue injury unfortunately also promotes cancer initiation. Identifying the possibility to mitigate cell plasticity negative effects is a key area of interest. The pancreas itself has a remarkable capacity for cell plasticity that promote cancer initiation (*1*). It is widely accepted that autophagy machinery, as part of the intracellular degradation system, plays a key role in the exocrine pancreas (*2, 3*), and is a critical component of the homeostasis control system for acinar cells, which have a remarkable capacity to synthesize secreted proteins. Autophagy is generally considered as a protective cellular process, as largely demonstrated by genetic disruption of macro-autophagy machinery encoding genes such as ATG7, ATG5 that, for instance, led to accumulation of precancer lesions (*3–9*). It is unclear whether and how these two protective mechanisms are integrated to maintain tissue homeostasis, and pancreatic acinar cells serve as a paradigmatic physiological model to evaluate the role of autophagy in cell plasticity.

Phosphoinositide 3-kinases (PI3Ks) are important lipid kinases producing D3-phosphorylated phosphoinositides that organize functional protein complexes regulating various biological processes. The sole class III PI3K, VPS34, is the most ancient of the eight vertebrate PI3K isoforms. It was initially identified in yeast and plays an essential role in vacuolar homeostasis (*10, 11*). The ubiquitous VPS34 is found in complexes with its regulatory protein kinase subunit, VPS15, and specifically catalyses the conversion of phosphatidylinositol into phosphatidylinositol-3-monophosphate (PI-3-P). In vertebrates, this lipid, present in relatively low amounts in cells, directly regulates endosomal trafficking, autophagosome formation, constantly controlling a basal flux of autophagy.

Autophagy has a dual role in cancer. Increased levels of autophagy promote growth of established tumours as well as treatment resistance at the progressive stage (e.g. in pancreatic cancer (*12*)). Reversely, tissular stress increases autophagy which is beneficial for healthy tissue repair. Increased levels of autophagy members promote the maintenance of homeostasis and the elimination of cellular damages thus preventing tumour initiation. As a consequence, a deficient level of autophagy machinery proteins promotes early cancer lesion development. During precancer inception, the autophagy machinery dysregulation is associated with the promotion of a selective immune environment that is permissive for cancer development (*13*); it leads to a selective metabolic niche that can be assessed through plasma measurements (*14, 15*) and is associated with most cancer risk factors (e.g. aging, diabetes, inflammatory disease) in Human (*16*).

We hypothesized and examined whether the class III PI3K VPS34 played a crucial role in the pancreas, providing a unique model to explore the importance of autophagy induction in regulating cell plasticity. Unexpectedly, our findings diverged from the sole autophagy machinery dysregulation, where autophagy impairment resulted in chronic inflammation or the development of precancerous lesions (*3–9*). Instead, we discovered a novel protective mechanism triggered from autophagy deregulation, which both prevents tissue damage and cell plasticity. This protective mechanism maintained acinar cells in a transcriptional state resistant to oncogene-induced cell plasticity (*1*), now enabling the possibility to prevent environmentally-induced increased cancer risk.

### VPS34 controls pancreatic homeostasis

We and others previously showed that pancreatic exocrine cells were plastic, transdifferentiating between acinar and ductal differentiation (*17*). Next, we measured exocrine cell autophagy capacity. A basal level of autophagy in murine ex-vivo primary culture of acini was measured, as lysozyme inhibitor Chloroquine (CQ) blocked autophagy flux and led to a significant two–fold increase of LC3-II (**Fig. 1A**). Interestingly, we identified a negative association between autophagic flux and VPS34 protein level (**Fig. 1A**) as shown by others(*18*). In Human healthy pancreas or in Human pancreas with chronic pancreatitis, VPS34^high^ positive staining was restricted to acinar cells; we observed a significant increase in population of VPS34^low^ acinar cells in pathological condition (ranging from moderately inflamed to severely inflamed acinar tissue), showing that low and high levels of VPS34 is found in acinar cells (**Fig.1B, Fig. S1)**. To understand the physiological role of decreased VPS34, we used a mouse model where exon 21 of the kinase domain of *PIK3C3* was flanked with loxP sites (VPS34lox/lox) to inactivate its activity in acinar cells (*19–21*) (**Fig. S2A**). The conditional and tamoxifen-inducible deletion in acinar cells, was obtained by breeding VPS34lox/lox mice with Elas-CreERT2 mice (*22*) to generate Elas-CreERT2+; VPS34lox/lox mice (referred as V34). Littermates not expressing Elas-CreERT2+ were used as controls and were treated with tamoxifen to account for possible action of tamoxifen injection alone. Full recombination was confirmed through genotyping of the pancreas (**Fig. S2B**). Pancreas were analysed at 2, 4, 8, 16 and 52 weeks post-recombination (**Fig. 1C**). The major unexpected phenotype was the absence of chronic pancreatitis (CP), as shown in other autophagic protein-deficient models (*3, 23*), even though we observed the presence of lipidic inclusions associated with mild signs of resolved resident inflammation (enlargement of the collagen fibres around the ducts) (**Fig. 1D** shows the phenotype compared to Human CP). Western blot analysis of whole pancreas lysates demonstrated that deletion of exon 21 in VPS34 did not reduce the stability of VPS34 nor of its complexes (**Fig. S2C**). Of note, V34 mice presented normal weight evolution, pancreas mass evolution and glycaemia levels, showing that the endocrine compartment of the pancreas was spared from VPS34 inactivation (**Fig. S3A**,**B**). Absence of CP phenotype was confirmed by unchanged levels of markers of exocrine cells, absence of increased amylase activity in the plasma and no significant changes in the number or diameter of duct-like structures at all ages tested (**Fig. S3C-E**).

**Fig. 1.**
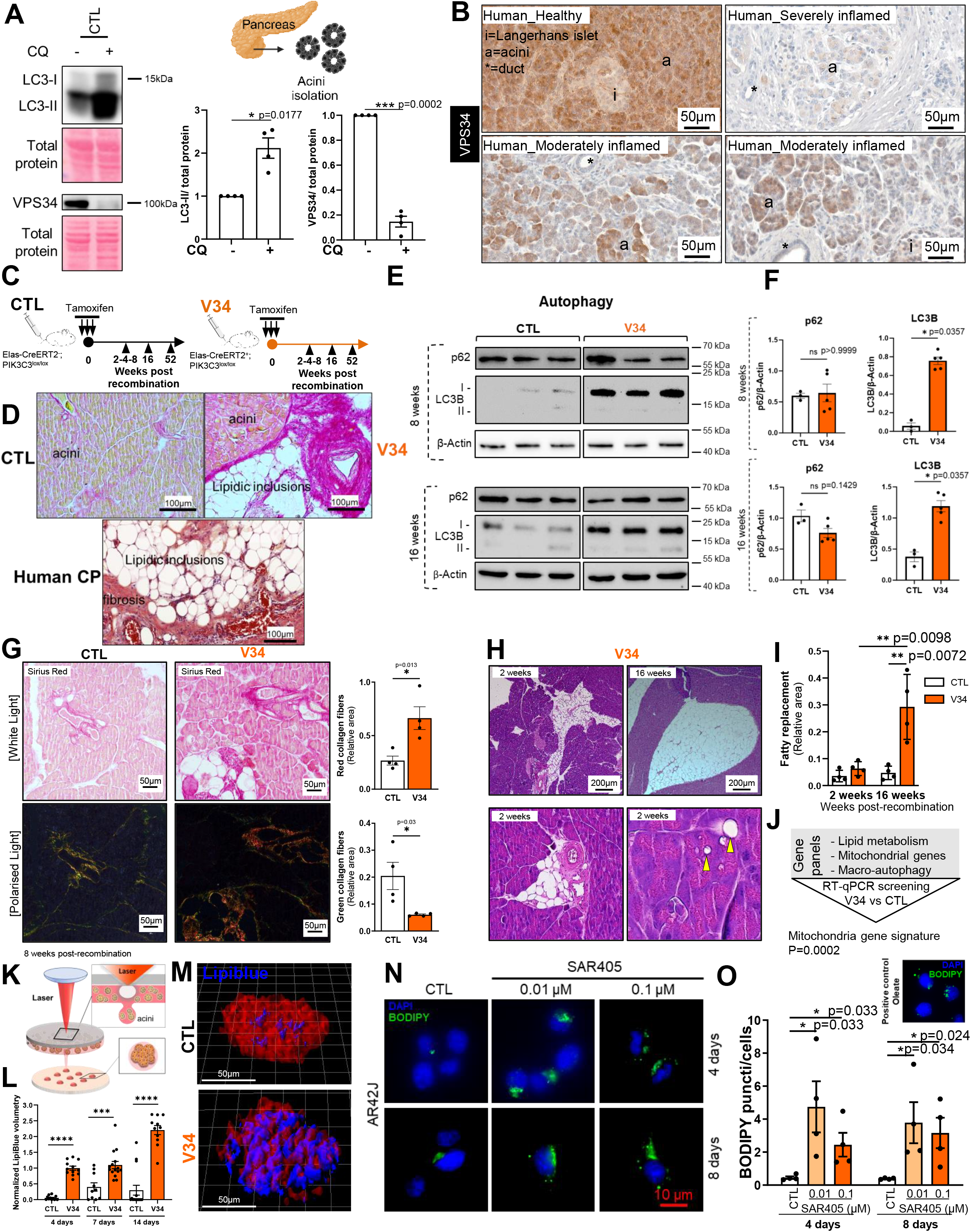
VPS34 controls pancreatic homeostasis, and its inactivation does not lead to chronic pancreatitis but to lipidic accumulation. (A) Western blot analysis showing VPS34 and the autophagosome marker LC3B on isolated acinar cells from CTL mice. Isolated primary acinar cells treated with or without CQ (10µM; 24h). Protein levels were normalized with total protein. Mean SEM, (*p<0.05, ** p<0.01, *** p<0.001), n = at least 4. (B), Immunohistochemistry of VPS34 in healthy and inflamed human acini; a highlights acini areas; i highlights Langerhans islets; asterisks highlights ducts. Scale as indicated. Clinical data and lower magnification images of the pancreas are shown in Fig. S1. N=8 patients. (C) Scheme of time points for evaluating the effects of VPS34 inactivation. Both Elas-CreERT2-control (CTL) and Elas-CreERT2+;PIK3C3^lox/lox^ (V34) mice were injected i.p. with tamoxifen as described. (D) Representative H&E staining of CTL and V34 52 week-old pancreas compared to human chronic pancreatitis. Scale as indicated. (E, F) Western blot analysis showing proteins involved in autophagy on pancreas tissues from CTL and V34 mice. Protein levels were normalized with β-Actin. Mean +/− SEM, (*p<0.05, Mann Whitney test), CTL n=3, V34 n=5. (G) CTL and V34 mice were injected by i.p. with tamoxifen as described above. Picrosirius red staining is performed on dissected pancreas and analysed with polarized light. Mean +/− SEM (*p<0.05, ** p<0.005), n=4 in each group. (H) Acinar cell vacuolation and fatty replacement in V34 mice pancreata 2 or 16 weeks after recombination. Yellow arrows show stressed acinar cells and inclusion of stressed vacuoles within pancreatic parenchyma only in V34 mice (n=4 in each group). (I) Fatty acid replacement was quantified. Mean +/− SEM (*p<0.05), n=4 in each group. (J) RT-qPCR on isolated acinar cells 52 weeks after tamoxifen injection, analysing the expression of signatures for lipid metabolism (8 genes), mitochondrial genes (9 genes) and macroautophagy (30 genes). Complete results are showed in Fig S3 A-E. (K) Bioprinting technology. Acini from CTL or V34 mice were bioprinted and the accumulation of lipids was followed using LipiBlue staining. (L) Quantification of normalized LipiBlue Volumetry in CTL and V34 acini cultured for 4, 7 and 14 days in the biofabricated exocrine pancreas. (n>10 in each group). (M) Representative acini in the biofabricated exocrine pancreas at 14d days; genotype is indicated. (N, O) Acinar cell line AR42J was treated with 0.01 or 0.1 µM of VPS34 inhibitor SAR405 and lipid accumulation was analysed using BODIPY staining and puncta/cell was quantified. Oleate treatment was used as a positive control. Mean +/− SEM (*p<0.05), n=4.

Next, we sought to establish a connection between the phenotypic alterations in V34 mice and the underlying mechanisms. VPS34 activity was shown to regulate endosomal trafficking and autophagosome formation (*10, 11*). We started by assessing changes at protein level of those cellular pathways that could be affected by VPS34 inactivation, namely autophagy and endocytic sorting. We evaluated the effects of VPS34 inhibition on autophagy by Western blot analyses of LC3B and SQSTM1/p62. At both 8- and 16-weeks post-recombination, we observed that LC3B and in particular LC3B-I accumulated in V34 mice, suggesting a possible alteration in autophagy at initiation and autophagosome formation step (**Fig. 1E,F**). Regarding the proteins involved in endocytosis, we observed that early-endosomal markers Rab5, and early-sorting endosome EEA1 levels were unchanged in V34 mice (**Fig. S2D**). The endosomal adaptor protein APPL1 level, also involved in endosome-dependent autophagy (*24*), was decreased at 16 weeks (**Fig. S2D**), indicating possible defects in endosome-regulated functions at a later stage.

Picro Sirius Red staining, used for analysing collagen fibre composition, showed mild signs of fibrosis in V34 mice (**Fig. 1G**). Quantification of red and green collagen fibres with polarized light revealed that at 8 weeks, the parenchyma of V34 mice was enriched in red fibres (type I collagen fibres) localised around ducts, in contrast to control mice which displayed more green fibres (type III collagen) (**Fig. 1G**). V34 mice also exhibited significant fatty replacement of pancreatic tissue. The fatty replacement was progressive: at the earliest point, the lipid accumulation occurred mainly in acinar cells around ducts, and then it replaced larger areas of the pancreas (**Fig. 1H,I**). Lipidic droplet accumulation is a common marker of autophagic flux defect shown to be associated with decreased mitochondrial β-oxidation (*25, 26*) and liver steatosis (*27*) as lipid droplet breakdown could be mediated by autophagy. A RT-qPCR screen in primary acini extracted from control and V34 mice showed a significant decrease of mitochondrial gene panel (**Fig. 1J**), as well as significant decrease of autophagy signal protein ULK2 mRNA and increase of the lipid storing protein ACACB mRNA in V34 mice compared to control mice (**Fig. S4A-E**). Part of acinar cells in V34 mice presented ultrastructural defects, with enlarged ER that resulted in increased ER-mitochondria contact coefficient (**Fig. S4F, top, S4G**), known to alter mitochondrial respiration efficiency (*26*). Transmission electron microscopy (TEM) also showed signs of mitochondrial fission, with smaller mitochondria embedded in ER (**Fig. S4F, bottom)**.

Presence of modified mitochondrial dynamics was confirmed by WB in tissue, with significant increased levels of p616DRP1 (**Fig. S4H**), a phosphorylated form of DRP1 that promotes fission (*28*). Pharmacological inactivation of VPS34 by SAR-405 and IN-1 (*29, 30*) in a pancreatic acinar cell line AR42J blocked nutrient starved EBSS-induced autophagy flux (restored levels of LC3I/II, p62), increased p616DRP1 level and tended to restore p616/p637DRP1 level (**Fig. S5A**,**B**).

The conservation of the pancreatic lobular structure and the presence of ducts within this new adipose tissue suggests an acinar origin for the adipocyte-like cells. To confirm this, we developed a method involving laser-assisted bioprinting to achieve the long-term culture of isolated acini (**Fig. 1K**). This bioprinted 3D primary culture of acini represents a significant and innovative advancement for modelling the exocrine pancreas (*31, 32*). CTL acini maintained their acinar cell morphology (**Fig. 1K**). In contrast, V34 acini progressively accumulated lipids, as evidenced by increased LipiBlue-positive staining over time, while CTL acini displayed only low baseline lipid staining with minimal changes over the same period (**Fig. 1L,M**). Finally, in AR42J acinar cells, pharmacological inhibition of VPS34 using SAR405 that blocked autophagy (**Fig. S5A**) led to significant accumulation of BOPIDY-positive puncta (**Fig. 1N,O**). Loss of VPS34 activity leads to possible transdifferentiation of some acinar cells into adipocyte-like cells.

### VPS34 activity controls acinar cell transcriptional identity

We next made the hypothesis that the heterogeneity of acinar cell VPS34 levels, indicative of heterogeneous autophagy flux levels observed in Human samples, could control acinar cell identity. To delve deeper into the mechanisms through which VPS34 loss was associated with maintenance of acinar cell function without inducing CP, we studied pancreatic cell population upon 4 to 8 weeks of VPS34 inactivation. We developed a protocol to isolate unique and viable pancreatic single cells followed by analysis by scRNAseq (**Fig. 2A, Fig. S6**) that identified 19 cell clusters belonging to the pancreatic epithelial lineage (acinar, acino-ductal, duct) or the microenvironment in both genetic background (**Fig. 2B-D, Fig. S7**). The most frequent acinar cell identity type found in CTL pancreas (Acinar_C0) (**Fig. 2E**) was characterized by mitochondrial gene expression (**Fig. 2F, Table S1**), in line with the result that V34 acinar cell expressed less mitochondrial genes (RT-qPCR, **Fig. S4E**). VPS34 inactivation altered the differentiation status of remaining acinar cells, leading to the emergence of acinar cell clusters named Acinar_C14,C15,C18 (**Fig. 2B,E**) that can be significantly differentiated from the most frequent cluster in basal condition by high levels of mRNAs of *REG* family (**Fig. 2F**). *<u>REG3A</u>* was highly increased in Acinar_C14,C15,C18 cells (**Fig. 2G**). In previously published sc-RNAseq data from Human pancreas, two acinar cell types were identified, *<u>REG3A</u>*-positive that accounted for around 5% of acinar cells vs. the remaining (undefined) acinar cells (*33*). The regenerating protein family, abbreviated as REG family are a group of small secretory proteins that are involved in the proliferation and differentiation of diverse cell types; they are important in protecting cells from death caused by damage or inflammation (*34, 35*). Using sc-RNAseq data from Human pancreas, we identified that *<u>REG3A</u>*-positive human acinar cells were associated with decreased levels of *GABARAPL2* and *MAP1LC3B* mRNAs, promoting macro-autophagy, and increased levels of lysosomal *CTSD* mRNA (**Fig. 2H**). In our model, C14, C15 and C18 *<u>REG</u>*-enriched acinar cell clusters were associated with a slight increase in innate immune system gene signatures (**Fig. 2I**). ScRNAseq also showed that VPS34 inactivation led to accumulation of mesenchymal and, endothelial cells, monocytes and macrophages (**Fig. 2B-D**), confirming the histological characterisation of reminiscent inflammation (**Fig. 1**). Similarly to Human data, C14, C15 and C18 *<u>REG</u>*-enriched acinar cell clusters were associated with a decrease in autophagy-related gene signatures, e.g. *Autophagosome vesicle trafficking lysosome, Non-conventional chaperone mediated autophagy, Conventional macroautophagy* (**Fig. 2I**). Hence, VPS34 activity controls acinar cell transcriptional identity.

**Fig. 2.**
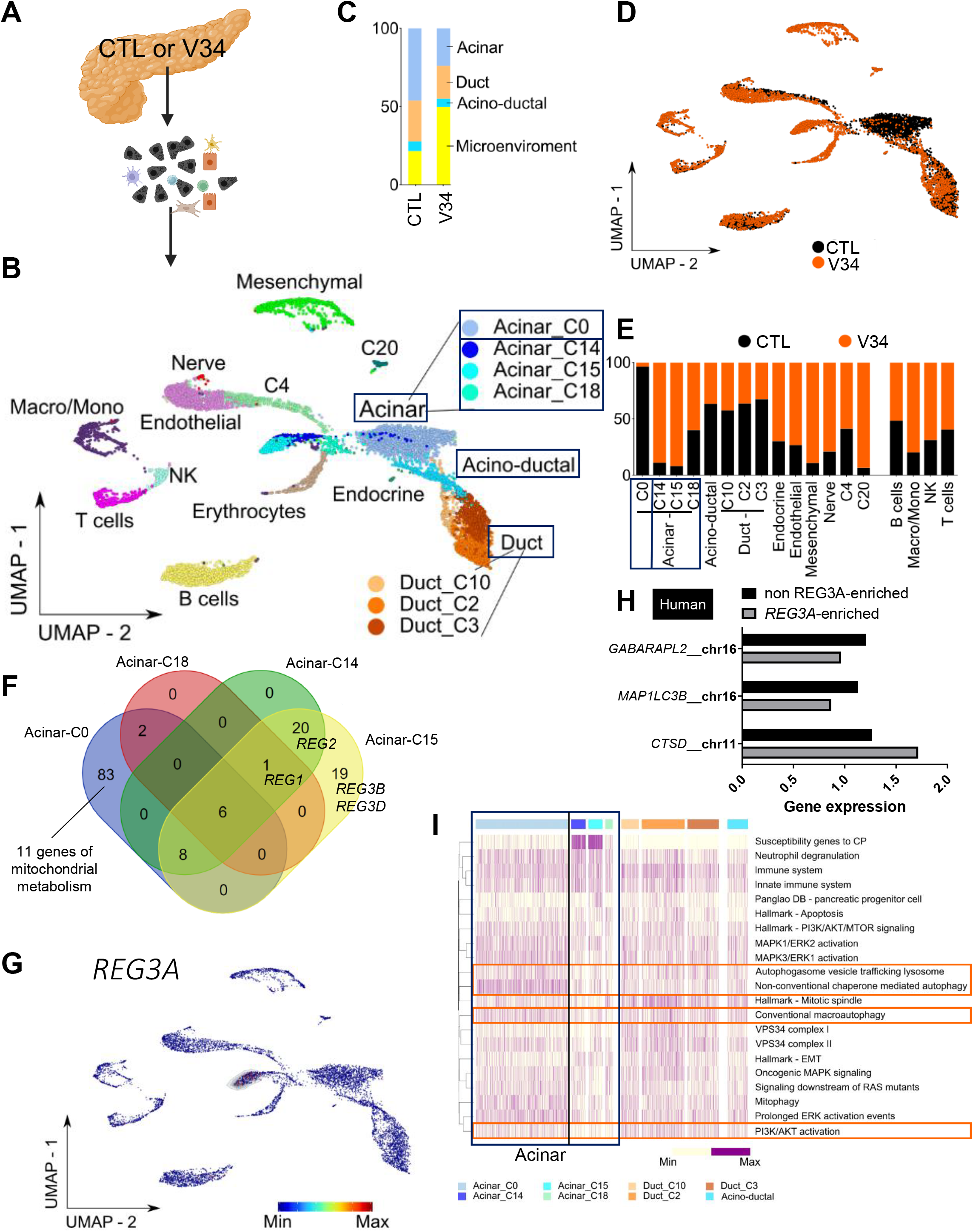
VPS34 activity controls acinar cell transcriptional identity. (**A**) Experimental design and sampling of 7 mice per genotypes in pools harvested at 4 and 8 weeks after tamoxifen injection (Control CTL: black, V34: orange). (**B**) Single-cell RNA sequencing (scRNAseq) Uniform Manifold Approximation and Projection (UMAP) of all cells per sample type. (**C**) Bar plot showing the proportion of the major three cell groups (Acinar, Acino-ductal and Duct) and the microenvironment of each sample type independently. (**D**) scRNAseq UMAP of all cells coloured by cell subset (legend). (**E**) Bar plot showing the proportion of each sample type among all the identified cell groups independently (same colour code as A). (**F**) Venn diagram showing the number of common differentially expressed genes (DEGs) between Acinar subsets (C0, C14, C15 and C18). (**G**) scRNAseq UMAP heatmap of REG3A gene expression among all cells, showing a strong association to Acinar C15 subset. (H) Bar plot of autophagy gene expression in human acinar cell clusters. Black: non-REG3A enriched, Grey: REG3A enriched. (I) Non-supervised hierarchical clustering heatmap of the enrichment scores of the most representative pathways among the main cell groups (Acinar, Acino-ductal and Duct).

### The newly identified acinar cell state present decreased REG3A protein level through an increase of the lysosomal compartment

Next, we aimed to molecularly characterize and functionally validate the transcriptomics analysis by analysing the Acinar_C14,C15,C18 cell clusters and Acinar_C0 cell cluster using in short term ex-vivo culture of V34 and CTL acinar cells, respectively. First, V34 acini presented significantly lower levels of REG3A protein (**Fig. 3A**) (also for REG2 but not of REG1, **Fig. S8**). In the Human *<u>REG3A</u>*-enriched acinar clusters, if autophagy *MAP1LC3* and *GABARAPL1* gene expression were decreased, *CTSD* encoding gene (lysosomal enzyme) expression was increased (**Fig. 2H**). Interestingly, lysosome inhibitor CQ partially rescued REG3A level only in V34 acini while the level of REG3A in CTL cells was not affected by CQ treatment (**Fig. 3A**), suggesting that REG3A is degraded by the lysosome compartment. Analysing the levels of autophagy markers in tissues is not sufficient to conclude about the autophagy status (*36*). To investigate further, we demonstrated that V34 acini had fewer

**Fig. 3.**
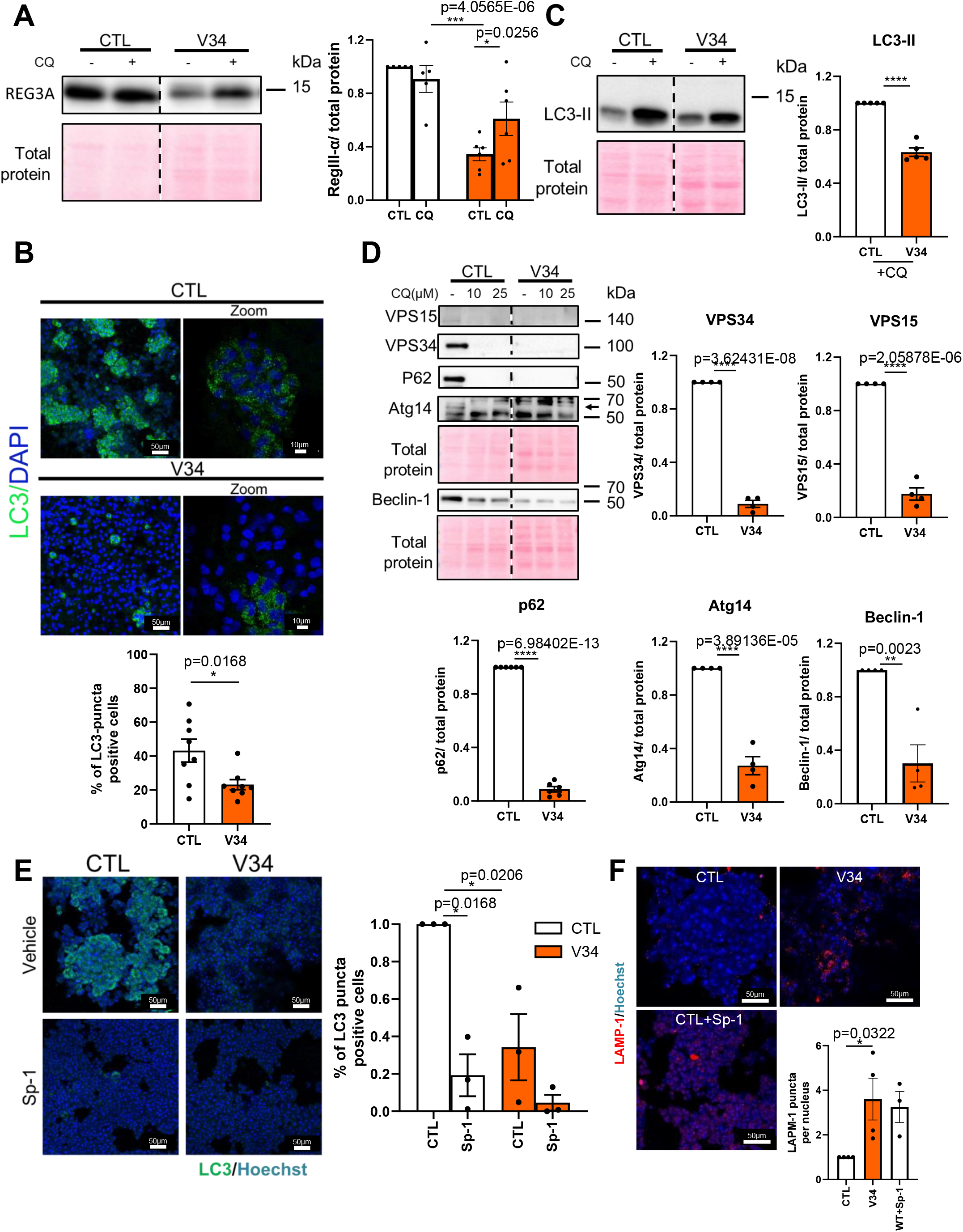
V34 acini enriched in acinar-C14,C15,C18 cell clusters present decreased REG3A protein level through an increase of the lysosomal compartment. (**A**) Western blot analysis showing REG3A on isolated acinar cells 5-9 weeks after tamoxifen injection from CTL or V34 mice. Isolated primary acinar cells treated with or without chloroquine CQ (10µM; 24 hrs). Protein levels were normalized with total protein. Mean +/− SEM, (*p<0.05, ** p<0.01, *** p<0.001), N=at least 5. (**B**) IF analysis of LC3B on isolated acinar cells 5-9 weeks after tamoxifen injection from CTL or V34 mice, 24 hours post culture. Scale bar= 50µM; Zoom=10µM. Mean +/− SEM, (*p<0.05, ** p<0.01, *** p<0.001), N=2. (**C**) Western blot analysis showing LC3B on isolated acinar cells 5-9 weeks after tamoxifen injection from CTL or V34 mice. Isolated primary acinar cells treated with or without chloroquine CQ (10µM; 24 hrs). Protein levels were normalized with total protein. Mean +/− SEM, (*p<0.05, ** p<0.01, *** p<0.001), N=at least 4. (**D**) Western blot analysis showing VPS15, VPS34, P62 and Atg14 and Beclin-1 on isolated acinar cells 5-9 weeks after tamoxifen injection from CTL or V34 mice. Isolated primary acinar cells treated with or without chloroquine CQ (10µM, 25µM; 24 hrs. Protein levels were normalized with total protein. Mean +/− SEM, (*p<0.05, ** p<0.01, *** p<0.001), N= 4 (**E**) IF analysis of LC3B on isolated acinar cells 5-9 weeks after tamoxifen injection from CTL or V34 mice, 24 hours post culture. Isolated primary acinar cells treated with or without spautin-1 [Sp-1] (2.5µM; 24 hrs). Scale bar= 50µM. Mean +/− SEM, (*p<0.05, ** p<0.01, *** p<0.001), N=3. (**F**) IF analysis of Lamp-1 on isolated acinar cells 5-9 weeks after tamoxifen injection from CTL or V34 mice, 24 hours post culture. Isolated primary acinar cells treated with or without spautin-1 [Sp-1] (2.5µM; 24 hrs). Scale bar= 50µM. Mean +/− SEM, (*p<0.05, ** p<0.01, *** p<0.001), N≥3.

LC3-positive puncta in short term ex-vivo culture compared to CTL acini suggesting that a lower number of V34 acini is performing autophagy (**Fig. 3B**). The decreased levels of accumulated LC3-II in V34 acini compared to CTL acini in the presence of CQ confirmed the low levels of autophagy flux in V34 acini that was suggested in Fig. 1A **(Fig. 3C)**. The levels of the autophagy receptor P62 and the proteins of complex 1, VPS34, VPS15, ATG14 and BECLIN-1 are controlled by the status of autophagy through the transcription factor TFEB(*18, 37*). The decrease in their basal levels in V34 acini compared to CTL acini further confirmed the lower activation of autophagy in V34 acini (**Fig. 3D**). CQ-induced inhibition in CTL acini showed a similar protein profile as V34 acini (**Fig. 3D**). Treatment with spautin (Sp-1) that promotes VPS34 complex I degradation (*via* Beclin-1 degradation (*38*) resulting in autophagy inhibition at early stages (*36*)) mimicked V34 phenotype of LC3-puncti decreased levels in CTL acini but did not have significant effect on V34 acini (**Fig. 3E**). Because decreased REG3A level in V34 acini was rescued by lysosome inhibitor chloroquine (CQ), suggesting that REG3A is degraded by the lysosome compartment, we analysed LAMP1 lysosome marker and found that V34 acini accumulated more LAMP1-positive cells. Spautin also mimicked V34 phenotype on LAMP-1-puncti accumulation CTL acini (**Fig. 3F**). These data shows that the V34 acini enriched in Acinar_C14,C15,C18 cell clusters present decreased REG protein levels (3A and 2), in particular REG3A through an increase of the lysosomal compartment. Importantly, REGs are known to be involved in promoting stemness and inflammation (*39*).

### VPS34 activity is required for acinar cell transformation

To functionally validate the importance of VPS34 in cell plasticity, we induced cell plasticity with oncogenic KRAS expression in CTL versus V34 background and analysed the evolution of the mouse cohort (Pdx1-Cre;VPS34+/+ vs Pdx1-Cre;VPS34^lox/lox^)(**Fig. 4A**). Oncogenic KRAS expression in pancreatic progenitor lead to a sequence of precancer lesion accumulation after initial acinar-to-ductal transdifferentiation (*17*). VPS34 inactivation fully protected from mutant KRAS induced lethality (**Fig. 4B**) and oncogenicity (**Fig. 4C**).

**Fig. 4.**
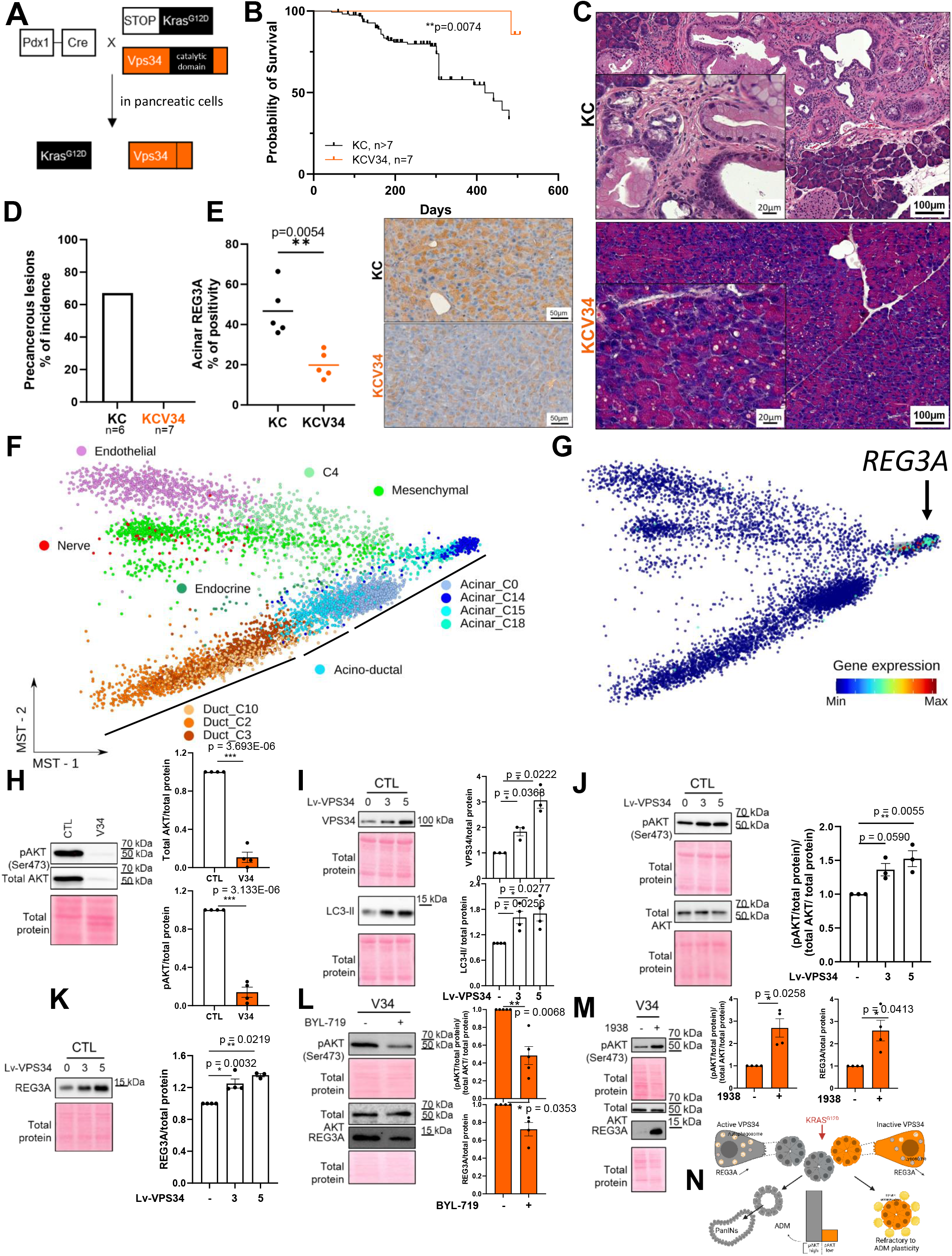
VPS34 activity is required for acinar cell transformation. **(A)** Gene construct of pancreatic-restricted KRAS mutated and VPS34 inactive mice (KCV34). In the gene PIK3C3 encoding VPS34, the kinase domain corresponds to exons 17-26. Deletion of exon 21 leads to the deletion from Ala730 to Thr754 in the translated protein. KRASGD12D expression from endogene locus is possible through the Lox-STOP-Lox cassette. STOP is a transcriptional Stop element. The recombination is mediated by Cre-recombinase under Pdx1 promoter control. (**B**) Survival curve of KC and KCV34 mice. Mantel-Cox statistical test, * p<0.05, ** p<0.01, *** p<0.001. N as indicated. (**C**) Representative HE staining of pancreas of KC and KCV34 mice showing precancerous lesions in KC and no lesion but vacuoles in acinar cells in KCV34. **(D)** Percentage of incidence of pancreatic precancerous lesions in KC and KCV34 mice. N as indicated. **(E)** Immunohistochemical staining of REG3A in pancreatic tissues from KC and KCV34 mice. Individual values and mean, Student t-test (* p<0.05, ** p<0.01, *** p<0.001), N=5 each group. (**F**) Representation of cell trajectory from scRNAseq data. (**G**) scRNAseq cell trajectory of *REG3A* gene expression among all cells, showing a strong association to Acinar C15 subset. (**H**) Western blot analysis showing pAKT (Ser473) and total AKT on isolated acinar cells 5-9 weeks after tamoxifen injection from CTL or V34 mice 24 hours post culture. Protein levels were normalized with total protein. Mean +/− SEM, (* p<0.05, ** p<0.01, *** p<0.001), N=4. (**I**,**J**,**K**) Western blot analysis showing (I) VPS34, LC3-II, (J) pAKT (Ser473), total AKT and (K) REG3A on isolated acinar cells 5-9 weeks after tamoxifen injection from CTL mice. Isolated primary acinar cells were transduced with polybrene or Lv-VPS34 (MOI= 3 or 5; 24 hours). Protein levels were normalized with total protein. Mean +/− SEM, (* p<0.05, ** p<0.01, *** p<0.001), N = at least 3. (**L**,**M**) Western blot analysis showing pAKT (Ser473), total AKT and REG3A on isolated acinar cells 5-9 weeks after tamoxifen injection from V34 mice. Isolated primary acinar cells were treated (L) without or with BYL-719 (1µM; 24h) or (M) without or with 1938 (10µM; 24h). Protein levels were normalized with total protein. Mean +/− SEM, (* p<0.05, ** p<0.01, *** p<0.001), N = at least 4. (**N**) Graphical abstract.

Strikingly, no precancer acinar-to-ductal ADM nor pancreatic intra-epithelial neoplasia PanIN were observed (**Fig. 4C**). Besides, the parenchyma was not replaced by large fatty deposits, only vacuoles in acinar cells were observed. The penetrance of the phenotype was 100% (**Fig. 4D**), hence, the phenotype was stronger than PI3Kα inactivation in oncogenic KRAS background, mouse model in which we and others previously observed some mice with little number of PanINs (*17, 40*). Acinar KCV34 cells presented decreased REG3A and REG1, but no change in Ki67 levels (**Fig. 4E, S9C**); REG2 levels were unchanged (**Fig. S9**). Trajectory analysis of scRNAseq data confirmed our results where we calculated that V34-enriched acinar clusters with high *REG3A* were exiting the acinar-to-ductal sequence at mRNA level (**Fig. 4F, G**). It is important to note that, *in vivo*, no spontaneous transdifferentiation in duct-like cells was observed in V34 mice at any observed time points (**Fig. 1C,D**), in contrast to other autophagy machinery-deficient models (*3, 4*). In AR42J acinar cell line, VPS34 pharmacological inhibition with SAR405 did not change acinar nor ductal gene expression (**Fig. S10A-D**) nor the number of cells with duct-like morphology (**Fig. S10E**) that was induced by the acinar-to-ductal (ADM) trans-differentiating Dexamethasone and Epithelial Growth Factor (EGF) cocktail (*17*).

We next sought to explain why C14, C15, C18 acinar cell clusters were refractory to cell plasticity. We first observed that these cells presented decreased *PI3K/AKT activation* gene signatures which are critical for epithelial cell plasticity (**Fig. 2I, 4G**) (*17*). We validated scRNAseq results at post-translational and protein level in V34 acini, showing a significant decreased pAKT/AKT ratio as well as AKT levels (**Fig. 4H**). To demonstrate direct causality between VPS34, control of autophagy and of REG3A and AKT phosphorylation levels in Acinar_C0 cell cluster, we transduced CTL acinar cell cultures with lentiviruses encoding for VPS34 (with *PIK3C3* cDNA). Overexpression of PIK3C3 led to VPS34 accumulation, increased LC3II levels indicating autophagy activation (**Fig. 4I**) and increased pAKT/AKT ratio, indicative of PI3K activation without increasing AKT levels (**Fig. 4J**). REG3A levels were also significantly increased (**Fig. 4K**). To demonstrate the role of PI3K blockage in controlling REG3A level in Acinar_C14,C15,C18 cell clusters, we searched and showed that PI3Kα pharmacological inhibition with BYL-719 decreased both pAKT/AKT ratio and REG3A in V34 acini (**Fig. 4L**). Lastly, to fully confirm causality between PI3Kα activation and REG3A increased level via AKT phosphorylation, we treated V34 acini with PI3Kα pharmacological activator 1938 compound (*41*) and observed an increased pAKT/AKT and REG3A levels, reversing V34 phenotype (**Fig. 4M**). VPS34^high^ cell state associated high VPS34 activity and the capability to perform macro-autophagy is necessary for cell plasticity and controls PI3Kα-dependent level of proinflammatory REG3A.

## Conclusions and discussion

In conclusion, VPS34 inactivation, instead of promoting acinar-to-ductal transdifferentiation, as observed in other stress or autophagy machinery defects, protected the pancreas from acinar-to-ductal plasticity, spontaneous chronic pancreatitis and formation of precancer lesions (**Fig. 4N, Graphical Abstract**).

Mechanistically, the differentiated acinar cells changed their transcriptional identity to a state less susceptible to cancer promotion by oncogenic KRAS, through reduced activation of classI PI3K and increased lysosomal degradation of the proinflammatory REG3A. Genetic disruption of macro-autophagy through autophagosome genes such as ATG7, ATG5 led to the discovery of their tumour suppressor role (*3–9*). We did not observe defects usually associated with ROS production and cell death (*3*) that was recently shown to be controlled by VPS34 in cancer progression models (*42*). Here, we find that VPS34 activity loss highlighted a novel protective mechanism arising from induced macro-autophagy deregulation and deregulated lysosomal compartment.

While the heterogeneity of the acinar cells was identified a while ago by novel scRNAseq method (*33, 43*), our study is one of the first to highlight the importance of a less frequent transcriptional state of acinar cells in the response to stress. This tissular response was unexpected. Differentiated cell plasticity plays a key role in tissue homeostasis in general (see other examples in (*44*), and this mechanism is largely co-opted by cancer initiating cells. Identifying ways of preventing acinar-to-ductal metaplasia makes possible intervention to block epithelial memory of inflammation, that was recently shown to promote cancer progression (*1*). Our work now enables strategies for protection from stress-induced cell plasticity and cancer initiation.

## Supporting information

all supplemental data and info

## Acknowledgments

We are grateful to SigDYN members including our talented visiting students, past and present, for their technical support, sample banks, common tools, scientific and protocol discussions, Prof. B Vanhaesebroeck and Dr B Bilanges (UCL, UK) for VPS34lox mice and feedback on VPS34 biochemistry, members of ART for discussion, Prof. A Couvelard, practioners and patients for clinical samples, UMS006/CREFRE, Anexplo Platform, Toulouse (mouse breeding & experimental zone; ENI core platform), the CRCT core technology platform (in particular Laetitia Ligat, Marie Tosolini, Fred Pont, Nathalie Saint-Laurent, Manon Farce, Tiphaine Fraineau, Loic Van Den Berghe), ImagIN platform (FX Fresnois), Histology platform (Christèle Ségura, I2MC, Inserm 1048, Toulouse, France), Dr S Severin (I2MC, Toulouse) for sharing tips for genotyping and mouse colonies and Dr J Laporte for access to IGBMC TEM.

## Funding

Agence National de la Recherche, Toucan, an integrated research program on Signal-targeted Drug Resistance, Laboratoire d’Excellence (JGG)

Fondation ARC, ARCPJA2021060003932 (JGG)

Fondation ARC, ARCPGA2022120005630_6362-3 (JGG)

Fondation ARC, 4^th^ Y PhD salary, (FRD, JGG)

Fondation ARC, 4^th^ Y PhD salary, (CC, JGG)

MSCA-ITN/ETN PhD-PI3K, Project ID: 675392, salary for FRD (JGG)

Agence National de la Recherche, ANR-JCJC-Radiance, including salary for BT (JGG) Cancéropôle GSO (JGG)

Ligue Nationale Contre le Cancer, salary for CC (JGG)

Institut National du Cancer INCa, PLBIO including salary of MdL, HS, INCa_19285 (JGG)

Fondation Toulouse Cancer Santé, Pacmine including salary for HS (JGG)

Hôpitaux de Toulouse, salary for CG (CG, JGG)

MSCA-Erasmus, bursary for AV (AV, JGG).

## Author contributions

Investigation: FRD, HS, CG, CH, PCA, RDA, NT, AV, CC, CV, ES, ND, MdL, CJ, HDO, MD, BT, JGG

Formal analysis: FRD, HS, CG, CH, PCA, AC, BT, JGG

Data curation: PCA, JGG

Methodology: FRD, HS, CG, CH, PCA, RDA, NT, CV, ES, CJ, HDO, MD, BT, JGG

Ressources: AC, VR, CJ, MD, JGG

Visualization: FRD, HS, CG, CH, PCA, AC, BT, JGG

Conceptualization: JGG

Validation: HDO, CJ, MD, BT, JGG

Funding acquisition: JGG

Project administration: JGG

Supervision: HDO, MD, BT, JGG

Writing – original draft: FRD, JGG

Writing – original draft of Material and Methods: FRD, HS, CG, CH, CV, PCA, BT, JGG

Writing – review & editing: all authors

## Data and materials availability

All data, code, and materials used in the analysis are available upon request. Accession number is GSE288860: the scRNADeq data studied time evolution of pancreatic cell population upon 4 to 8 weeks of VPS34 inactivation. A protocol to isolate unique and viable pancreatic single cells followed by analysis by scRNAseq that identified 19 cell clusters belonging to the pancreatic epithelial lineage (acinar, acino-ductal, duct) or the microenvironment in both genetic background was developed.

## Supplementary Materials

Materials and Methods

Supplementary Text

Figs. S1 to S10

## References and Notes

1. E. Del Poggetto, I.-L. Ho, C. Balestrieri, E.-Y. Yen, S. Zhang, F. Citron, R. Shah, D. Corti, G. R. Diaferia, C.-Y. Li, S. Loponte, F. Carbone, Y. Hayakawa, G. Valenti, S. Jiang, L. Sapio, H. Jiang, P. Dey, S. Gao, A. K. Deem, S. Rose-John, W. Yao, H. Ying, A. D. Rhim, G. Genovese, T. P. Heffernan, A. Maitra, T. C. Wang, L. Wang, G. F. Draetta, A. Carugo, G. Natoli, A. Viale, Epithelial memory of inflammation limits tissue damage while promoting pancreatic tumorigenesis. Science 373, eabj0486 (2021).

2. K. L. Bryant, C. A. Stalnecker, D. Zeitouni, J. E. Klomp, S. Peng, A. P. Tikunov, V. Gunda, M. Pierobon, A. M. Waters, S. D. George, G. Tomar, B. Papke, G. A. Hobbs, L. Yan, T. K. Hayes, J. N. Diehl, G. D. Goode, N. V. Chaika, Y. Wang, G.-F. Zhang, A. K. Witkiewicz, E. S. Knudsen, E. F. Petricoin, P. K. Singh, J. M. Macdonald, N. L. Tran, C. A. Lyssiotis, H. Ying, A. C. Kimmelman, A. D. Cox, C. J. Der, Combination of ERK and autophagy inhibition as a treatment approach for pancreatic cancer. Nature Medicine 25, 628–640 (2019).

3. K. N. Diakopoulos, M. Lesina, S. Wörmann, L. Song, M. Aichler, L. Schild, A. Artati, W. Römisch-Margl, T. Wartmann, R. Fischer, Y. Kabiri, H. Zischka, W. Halangk, I. E. Demir, C. Pilsak, A. Walch, C. S. Mantzoros, J. M. Steiner, M. Erkan, R. M. Schmid, H. Witt, J. Adamski, H. Algül, Impaired autophagy induces chronic atrophic pancreatitis in mice via sex- and nutrition-dependent processes. Gastroenterology 148, 626–638.e17 (2015).

4. K. Görgülü, K. N. Diakopoulos, J. Ai, B. Schoeps, D. Kabacaoglu, A.-F. Karpathaki, K. J. Ciecielski, E. Kaya-Aksoy, D. A. Ruess, A. Berninger, M. Kowalska, M. Stevanovic, S. M. Wörmann, T. Wartmann, Y. Zhao, W. Halangk, S. Voronina, A. Tepikin, A. M. Schlitter, K. Steiger, A. Artati, J. Adamski, M. Aichler, A. Walch, M. Jastroch, G. Hartleben, C. S. Mantzoros, W. Weichert, R. M. Schmid, S. Herzig, A. Krüger, B. Sainz, M. Lesina, H. Algül, Levels of the Autophagy-Related 5 Protein Affect Progression and Metastasis of Pancreatic Tumors in Mice. Gastroenterology 156, 203–217.e20 (2019).

5. A. Takamura, M. Komatsu, T. Hara, A. Sakamoto, C. Kishi, S. Waguri, Y. Eishi, O. Hino, K. Tanaka, N. Mizushima, Autophagy-deficient mice develop multiple liver tumors. Genes Dev. 25, 795–800 (2011).

6. J. Y. Guo, B. Xia, E. White, Autophagy-mediated tumor promotion. Cell 155, 1216–1219 (2013).

7. A. Yang, G. Herter-Sprie, H. Zhang, E. Y. Lin, D. Biancur, X. Wang, J. Deng, J. Hai, S. Yang, K.-K. Wong, A. C. Kimmelman, Autophagy Sustains Pancreatic Cancer Growth through Both Cell-Autonomous and Nonautonomous Mechanisms. Cancer Discovery 8, 276–287 (2018).

8. K. Yamamoto, A. Venida, J. Yano, D. E. Biancur, M. Kakiuchi, S. Gupta, A. S. W. Sohn, S. Mukhopadhyay, E. Y. Lin, S. J. Parker, R. S. Banh, J. A. Paulo, K. W. Wen, J. Debnath, G. E. Kim, J. D. Mancias, D. T. Fearon, R. M. Perera, A. C. Kimmelman, Autophagy promotes immune evasion of pancreatic cancer by degrading MHC-I. Nature 581, 100–105 (2020).

9. J. Y. Guo, E. White, Role of Tumor Cell Intrinsic and Host Autophagy in Cancer. Cold Spring Harb Perspect Med 14, a041539 (2024).

10. B. Vanhaesebroeck, J. Guillermet-Guibert, M. Graupera, B. Bilanges, The emerging mechanisms of isoform-specific PI3K signalling. Nat. Rev. Mol. Cell Biol. 11, 329–341 (2010).

11. B. Bilanges, Y. Posor, B. Vanhaesebroeck, PI3K isoforms in cell signalling and vesicle trafficking. Nat. Rev. Mol. Cell Biol. 20, 515–534 (2019).

12. K. L. Bryant, C. A. Stalnecker, D. Zeitouni, J. E. Klomp, S. Peng, A. P. Tikunov, V. Gunda, M. Pierobon, A. M. Waters, S. D. George, G. Tomar, B. Papke, G. A. Hobbs, L. Yan, T. K. Hayes, J. N. Diehl, G. D. Goode, N. V. Chaika, Y. Wang, G.-F. Zhang, A. K. Witkiewicz, E. S. Knudsen, E. F. Petricoin, P. K. Singh, J. M. Macdonald, N. L. Tran, C. A. Lyssiotis, H. Ying, A. C. Kimmelman, A. D. Cox, C. J. Der, Combination of ERK and autophagy inhibition as a treatment approach for pancreatic cancer. Nature Medicine 25, 628–640 (2019).

13. S. Rao, L. Tortola, T. Perlot, G. Wirnsberger, M. Novatchkova, R. Nitsch, P. Sykacek, L. Frank, D. Schramek, V. Komnenovic, V. Sigl, K. Aumayr, G. Schmauss, N. Fellner, S. Handschuh, M. Glösmann, P. Pasierbek, M. Schlederer, G. P. Resch, Y. Ma, H. Yang, H. Popper, L. Kenner, G. Kroemer, J. M. Penninger, A dual role for autophagy in a murine model of lung cancer. Nat Commun 5, 3056 (2014).

14. E. White, J. M. Mehnert, C. S. Chan, Autophagy, Metabolism, and Cancer. Clin Cancer Res 21, 5037–5046 (2015).

15. A. J. Clarke, A. K. Simon, Autophagy in the renewal, differentiation and homeostasis of immune cells. Nat Rev Immunol 19, 170–183 (2019).

16. H.-Y. Chen, E. White, Role of autophagy in cancer prevention. Cancer Prev Res (Phila) 4, 973–983 (2011).

17. R. Baer, C. Cintas, M. Dufresne, S. Cassant-Sourdy, N. Schönhuber, L. Planque, H. Lulka, B. Couderc, C. Bousquet, B. Garmy-Susini, B. Vanhaesebroeck, S. Pyronnet, D. Saur, J. Guillermet-Guibert, Pancreatic cell plasticity and cancer initiation induced by oncogenic Kras is completely dependent on wild-type PI 3-kinase p110α. Genes Dev. 28, 2621–2635 (2014).

18. H. Tao, P. G. Yancey, J. L. Blakemore, Y. Zhang, L. Ding, W. G. Jerome, J. D. Brown, K. C. Vickers, M. F. Linton, Macrophage SR-BI modulates autophagy via VPS34 complex and PPARα transcription of Tfeb in atherosclerosis. Journal of Clinical Investigation 131, e94229 (2021).

19. C. Valet, M. Levade, G. Chicanne, B. Bilanges, C. Cabou, J. Viaud, M.-P. Gratacap, F. Gaits-Iacovoni, B. Vanhaesebroeck, B. Payrastre, S. Severin, A dual role for the class III PI3K, Vps34, in platelet production and thrombus growth. Blood 130, 2032–2042 (2017).

20. C. J. F. Courreges, E. C. M. Davenport, B. Bilanges, E. Rebollo-Gomez, J. Hukelmann, P. Schoenfelder, J. R. Edgar, D. Sansom, C. Scudamore, R. Roychudhuri, O. A. Garden, B. Vanhaesebroeck, K. Okkenhaug, Lack of phosphatidylinositol 3-kinase VPS34 in regulatory T cells leads to a fatal lymphoproliferative disorder without affecting their development. [Preprint] (2024). 10.1101/2024.01.08.574346.

21. C. J. F. Courreges, E. C. M. Davenport, B. Bilanges, E. Rebollo-Gomez, J. Hukelmann, P. Schoenfelder, J. R. Edgar, D. Sansom, C. L. Scudamore, R. Roychoudhuri, O. A. Garden, B. Vanhaesebroeck, K. Okkenhaug, Lack of phosphatidylinositol 3-kinase VPS34 in regulatory T cells leads to a fatal lymphoproliferative disorder without affecting their development. Front Immunol 15, 1374621 (2024).

22. A. Grimont, A. V. Pinho, M. J. Cowley, C. Augereau, A. Mawson, M. Giry-Laterrière, G. Van den Steen, N. Waddell, M. Pajic, C. Sempoux, J. Wu, S. M. Grimmond, A. V. Biankin, F. P. Lemaigre, I. Rooman, P. Jacquemin, SOX9 regulates ERBB signalling in pancreatic cancer development. Gut 64, 1790–1799 (2015).

23. L. Antonucci, J. B. Fagman, J. Y. Kim, J. Todoric, I. Gukovsky, M. Mackey, M. H. Ellisman, M. Karin, Basal autophagy maintains pancreatic acinar cell homeostasis and protein synthesis and prevents ER stress. Proc. Natl. Acad. Sci. U.S.A. 112, E6166–6174 (2015).

24. K. K. L. Wu, K. Long, H. Lin, P. M. F. Siu, R. L. C. Hoo, D. Ye, A. Xu, K. K. Y. Cheng, The APPL1-Rab5 axis restricts NLRP3 inflammasome activation through early endosomal-dependent mitophagy in macrophages. Nat Commun 12, 6637 (2021).

25. B. Bilanges, S. Alliouachene, W. Pearce, D. Morelli, G. Szabadkai, Y.-L. Chung, G. Chicanne, C. Valet, J. M. Hill, P. J. Voshol, L. Collinson, C. Peddie, K. Ali, E. Ghazaly, V. Rajeeve, G. Trichas, S. Srinivas, C. Chaussade, R. S. Salamon, J. M. Backer, C. L. Scudamore, M. A. Whitehead, E. P. Keaney, L. O. Murphy, R. K. Semple, B. Payrastre, S. Tooze, B. Vanhaesebroeck, Vps34 PI 3-kinase inactivation enhances insulin sensitivity through reprogramming of mitochondrial metabolism. Nat Commun 8, 1804 (2017).

26. C. Bosc, N. Broin, M. Fanjul, E. Saland, T. Farge, C. Courdy, A. Batut, R. Masoud, C. Larrue, S. Skuli, N. Espagnolle, J.-C. Pagès, A. Carrier, F. Bost, J. Bertrand-Michel, J. Tamburini, C. Récher, S. Bertoli, V. Mansat-De Mas, S. Manenti, J.-E. Sarry, C. Joffre, Autophagy regulates fatty acid availability for oxidative phosphorylation through mitochondria-endoplasmic reticulum contact sites. Nat Commun 11, 4056 (2020).

27. R. Singh, S. Kaushik, Y. Wang, Y. Xiang, I. Novak, M. Komatsu, K. Tanaka, A. M. Cuervo, M. J. Czaja, Autophagy regulates lipid metabolism. Nature 458, 1131–1135 (2009).

28. W. Chen, H. Zhao, Y. Li, Mitochondrial dynamics in health and disease: mechanisms and potential targets. Sig Transduct Target Ther 8, 333 (2023).

29. R. Bago, N. Malik, M. J. Munson, A. R. Prescott, P. Davies, E. Sommer, N. Shpiro, R. Ward, D. Cross, I. G. Ganley, D. R. Alessi, Characterization of VPS34-IN1, a selective inhibitor of Vps34, reveals that the phosphatidylinositol 3-phosphate-binding SGK3 protein kinase is a downstream target of class III phosphoinositide 3-kinase. Biochemical Journal 463, 413–427 (2014).

30. B. Ronan, O. Flamand, L. Vescovi, C. Dureuil, L. Durand, F. Fassy, M.-F. Bachelot, A. Lamberton, M. Mathieu, T. Bertrand, J.-P. Marquette, Y. El-Ahmad, B. Filoche-Romme, L. Schio, C. Garcia-Echeverria, H. Goulaouic, B. Pasquier, A highly potent and selective Vps34 inhibitor alters vesicle trafficking and autophagy. Nat Chem Biol 10, 1013–1019 (2014).

31. D. Hakobyan, C. Médina, N. Dusserre, M.-L. Stachowicz, C. Handschin, J.-C. Fricain, J. Guillermet-Guibert, H. Oliveira, Laser-assisted 3D bioprinting of exocrine pancreas spheroid models for cancer initiation study. Biofabrication 12, 035001 (2020).

32. C. Handschin, H. Shalhoub, A. Mazet, C. Guyon, N. Dusserre, E. Boutet-Robinet, H. Oliveira, J. Guillermet-Guibert, Biotechnological advances in 3D modeling of cancer initiation. Examples from pancreatic cancer research and beyond. Biofabrication 17 (2025).

33. M. J. Muraro, G. Dharmadhikari, D. Grün, N. Groen, T. Dielen, E. Jansen, L. van Gurp, M. Engelse, F. Carlotti, E. J. P. de Koning, A. van Oudenaarden, A Single-Cell Transcriptome Atlas of the Human Pancreas. Cell Systems 3, 385–394.e3 (2016).

34. C. Sun, X. Wang, Y. Hui, H. Fukui, B. Wang, H. Miwa, The Potential Role of REG Family Proteins in Inflammatory and Inflammation-Associated Diseases of the Gastrointestinal Tract. IJMS 22, 7196 (2021).

35. L. Wang, Y. Quan, Y. Zhu, X. Xie, Z. Wang, L. Wang, X. Wei, F. Che, The regenerating protein 3A: a crucial molecular with dual roles in cancer. Mol Biol Rep 49, 1491–1500 (2022).

36. H. Shalhoub, P. Gonzalez, A. Dos Santos, J. Guillermet-Guibert, N. Moniaux, N. Dupont, J. Faivre, Simultaneous activation and blockade of autophagy to fight hepatocellular carcinoma. Autophagy Reports 3, 2326241 (2024).

37. A. Bahrami, V. Bianconi, M. Pirro, H. M. Orafai, A. Sahebkar, The role of TFEB in tumor cell autophagy: Diagnostic and therapeutic opportunities. Life Sciences 244, 117341 (2020).

38. J. Liu, H. Xia, M. Kim, L. Xu, Y. Li, L. Zhang, Y. Cai, H. V. Norberg, T. Zhang, T. Furuya, M. Jin, Z. Zhu, H. Wang, J. Yu, Y. Li, Y. Hao, A. Choi, H. Ke, D. Ma, J. Yuan, Beclin1 Controls the Levels of p53 by Regulating the Deubiquitination Activity of USP10 and USP13. Cell 147, 223–234 (2011).

39. Q. Li, H. Wang, G. Zogopoulos, Q. Shao, K. Dong, F. Lv, K. Nwilati, X.-Y. Gui, A. Cuggia, J.-L. Liu, Z.-H. Gao, Reg proteins promote acinar-to-ductal metaplasia and act as novel diagnostic and prognostic markers in pancreatic ductal adenocarcinoma. Oncotarget 7, 77838–77853 (2016).

40. C.-Y. C. Wu, E. S. Carpenter, K. K. Takeuchi, C. J. Halbrook, L. V. Peverley, H. Bien, J. C. Hall, K. E. DelGiorno, D. Pal, Y. Song, C. Shi, R. Z. Lin, H. C. Crawford, PI3K regulation of RAC1 is required for KRAS-induced pancreatic tumorigenesis in mice. Gastroenterology 147, 1405–1416.e7 (2014).

41. G. Q. Gong, B. Bilanges, B. Allsop, G. R. Masson, V. Roberton, T. Askwith, S. Oxenford, R. R. Madsen, S. E. Conduit, D. Bellini, M. Fitzek, M. Collier, O. Najam, Z. He, B. Wahab, S. H. McLaughlin, A. W. E. Chan, I. Feierberg, A. Madin, D. Morelli, A. Bhamra, V. Vinciauskaite, K. E. Anderson, S. Surinova, N. Pinotsis, E. Lopez-Guadamillas, M. Wilcox, A. Hooper, C. Patel, M. A. Whitehead, T. D. Bunney, L. R. Stephens, P. T. Hawkins, M. Katan, D. M. Yellon, S. M. Davidson, D. M. Smith, J. B. Phillips, R. Angell, R. L. Williams, B. Vanhaesebroeck, A small-molecule PI3Kα activator for cardioprotection and neuroregeneration. Nature 618, 159–168 (2023).

42. M. M. Swamynathan, S. Kuang, K. E. Watrud, M. R. Doherty, C. Gineste, G. Mathew, G. Q. Gong, H. Cox, E. Cheng, D. Reiss, J. Kendall, D. Ghosh, C. R. Reczek, X. Zhao, T. Herzka, S. Špokaitė, A. N. Dessus, S. T. Kim, O. Klingbeil, J. Liu, D. G. Nowak, H. Alsudani, T.-L. Wee, Y. Park, F. Minicozzi, K. Rivera, A. S. Almeida, K. Chang, R. P. Chakrabarty, J. E. Wilkinson, P. A. Gimotty, S. D. Diermeier, M. Egeblad, C. R. Vakoc, J. W. Locasale, N. S. Chandel, T. Janowitz, J. B. Hicks, M. Wigler, D. J. Pappin, R. L. Williams, P. Cifani, D. A. Tuveson, J. Laporte, L. C. Trotman, Dietary pro-oxidant therapy by a vitamin K precursor targets PI 3-kinase VPS34 function. Science 386, eadk9167 (2024).

43. D. Wollny, S. Zhao, I. Everlien, X. Lun, J. Brunken, D. Brüne, F. Ziebell, I. Tabansky, W. Weichert, A. Marciniak-Czochra, A. Martin-Villalba, Single-Cell Analysis Uncovers Clonal Acinar Cell Heterogeneity in the Adult Pancreas. Dev. Cell 39, 289–301 (2016).

44. A. J. Merrell, B. Z. Stanger, Adult cell plasticity in vivo: de-differentiation and transdifferentiation are back in style. Nat Rev Mol Cell Biol 17, 413–425 (2016).

