## Supplementary material for "Autophagy driven by VPS34 enables differentiated cell plasticity and cancer initiation": all supplemental data and info

**The PDF file includes:**

Materials and Methods
Figs. S1 to S10
References

**Other Supplementary Materials for this manuscript include the following:**

MTA for Class III mice

### Materials and Methods

Reagents

Chloroquine (C6628, Sigma-Aldrich) was used to block autophagy. Spautin-1 (SML0440, Sigma-Aldrich /Merk) was used to inhibit the early steps of autophagy. UCL-TRO-1938 (HY-154848, MedChemExpress) was used to activate PI3Kα. BYL-719 (Alpelisib)(A-4477, MedChemExpress) was used to inhibit PI3Kα. SAR405 (A8883, ApexBio Technology) and Vps34-IN1 (B6179, ApexBio Technology) were used to inhibit VPS34. All drugs were dissolved in DMSO (UD8050-05-C, Euromedex). All used reagents and references are listed in Table S1.

Human samples

Human pancreatitis sample were collected according French and European legislation, with informed consent obtained after the nature and possible consequences of the studies were explained. Pancreatic samples were retrospectively retrieved from the Pathology Department of Beaujon Hospital, Clichy. (IRB: 2018-A02712-53). One paraffin-embedded block of pancreatitis was selected in each case. Five 4 µm sections were performed on each block, for hematoxylin-eosin.

Mice

VPS34 was inactivated in pancreatic acinar cells (recombination of the loxed PIK3C3 locus (*1*) encoding for VPS34 by active CreER expressed in Elastase-positive cells (*2*, *3*) under tamoxifen treatment (*4*). In details, VPS34lox/lox;Elas-CreERT2+ mice (V34) on C57/B6J background were generated by breeding. Controls are VPS34lox/lox mice also subjected to tamoxifen treatment. Cre recombinase expression was induced in 4-week-old female and male mice by intraperitoneal injection of tamoxifen (1mg/15g body weight, T5648, Sigma) in corn oil (C8267, Sigma) for 3 consecutive days; this treatment was also used in the control cohort to account for the possible action of tamoxifen alone. Cohorts of 5 animals in each sex was done to account for possible sex bias.  LSL-KrasG12D knock-in from D. Tuveson, Mouse Models of Human Cancers Consortium repository, National Cancer Institute-Frederick) were crossbred with either only Pdx1-Cre from D.A. Melton, Harvard University, Cambridge, MA or with Pdx1-Cre and VPS34lox/lox from (*1*) on mixed background C57/B67 to obtain LSL-KrasG12D; Pdx1-Cre; VPS34lox/lox (named KCV34) and LSL-KrasG12D; Pdx1-Cre (named KC). Mice were followed (weight, imaging with Aixplorer, plasma, endpoints). All procedures and animal housing were performed according to the Institutional guidelines and Regulatory Bodies standards and those were approved by the ethical committee according to European legislation translated to French law as Décret 2013-118  February 1st, 2013, and certified by an authorization of experimentation delivered by the French Ministry of Higher Education and Research (MESR).

Tissue collection

Parts of the tissues (pancreas, liver, spleen, lung) were excised, then snap frozen in liquid nitrogen and stored at -80ºC until further use. Remaining parts were fixed for histological and immunostaining analysis according to each specific technique. Organs were weighted.

Plasma collection

Blood was collected by intracardiac puncture at sacrifice and then transferred to Eppendorf tubes containing 20µL 0.5M EDTA. Plasma was separated by centrifugation at 4000 rpm for 20 min at 4ºC and then stored at -80ºC until further use.

Amylase test

The amylase activity was determined using the Phadebas® Amylase Test. A modification was performed to the manufacture's kit to adjust the required sample volume to 10 µL. A "master mix" was prepared by pre-incubating 4 mL of distilled water at 37ºC for 5 min. A tablet was added to the pre-incubated water and then incubated for another 5 min at 37ºC. 10 µL of sample were pipetted into a 96-well plate. 200 µL of the master mix were added to each well. The plate was incubated at 37ºC for 15 min, then it was briefly stirred, and it was then centrifuged for 5 min at 4000 rpm. 100 µL of the supernatant was transferred to a new plate. The absorbance was measured at 620 nm against distilled water. All samples were performed in duplicates. The samples were prepared by diluting 10 µL of blood plasma in 190 µL mQ H2O. Saliva was used as a positive control in a 1/200 dilution in mQ H2O.

Genotyping and recombination

Genotyping was performed on the phalanx of 6-days-old mice. Recombination was performed on a part of snap frozen pancreas collected. DNA extraction was performed with a Sigma kit (Extraction solution: E7526, Tissue Preparation: T3073, Neutralization solution: N3910). Tissues are incubated 15 minutes with a mix of 100µL Extraction and 25µL Tissue preparation solutions by sample, followed by a DNA denaturation at 95°C during 5 minutes. The reaction is stopped by 100µL Neutralization Solution. Samples are centrifugated at room temperature during 5 minutes at 18 400g. Supernatant, containing DNA, is extracted, and stored at -20°C until further use. DNA solution is diluted to the tenth with a solution composed by specific forward primer and reverse primer (10µM) according to the gene of interest, water, and a mix containing the TaqPolymerase (RedExtract- N-Amp PCR Ready Mix, Sigma Ref R4775-12ML). ChemiDoc analysis required 2% agarose (Sigma) gel and 1Kb DNA ladder kit (Promega G5711). After the migration, gel was stained by SYBR® Safe solution (Invitrogen S33102).

Electron microscopy and ER-mitochondria contact coefficient (ERMICC) calculation

Pancreata sections of 1-2 mm3 were immersion-fixed using 2.5% (v/v) glutaraldehyde in 0.1 M cacodylate buffer for 4h at 4ºC. Sections were post-fixed in an aqueous solution 2% (w/v) OsO4 in 0.2 M cacodylate buffer for 2 h at 4ºC and then washed in 0.1 M cacodylate buffer. Tissue sections were dehydrated in graded ethanol, then in propylene oxide and embedded in Epon resin. Resin blocks were cured for 48 h at 60ºC sectioned (70 nm nominal) and placed on rhodium 400 mesh grids. Grids were contrasted with 3% uranyl acetate (2 h) followed by lead citrate (5 min). Grids were examined using Morgani 268 Transmission Electron Microscope (Philips/FEI) operating at 100kV, at Centre d'imagerie from the IGBMC in Strasbourg, France, and METI in CBI, Toulouse, France.

Endoplasmic reticulum (ER) and mitochondria areas were manually delineated on electron microscopy using the TrakEM2 pluging on ImageJ. Mitochondria were circled by a perimeter of 30 nm corresponding to the theoritical maximum distance separating ER and mitochondria in the MAM (mitochondria-associated membranes). We quantified each portion of ER comprised inside this perimeter and the mean ER-mitochondria distance for each ER portion measured. k

ER lengths (l) and k ER-mitochondria corresponding distances (d) were obtained for each mitochondria. The total ER length L was obtained by adding all l lengths. The following formula (adapted from (*5*)) was used to calculate the D mean ER-mitochondria distance over the whole mitochondria:

$$D = \frac{\sum_{i=0}^{i=k}(d_i * l_i)}{L}$$

The ER-mitochondria contact coefficient (ERMICC) was calculated with the following formula with $P$ being the perimeter of the mitochondria:

$$ERMICC = \frac{L}{P * D}$$

Bioprinting

Laser-assisted bioprinting was used to deposit droplets of extracted acini (as detailed below) on a methacrylated gelatin (5% w/v) biopaper. Extracted acini were suspended at 2.5 million/mL in methacrylated gelatin (GelMA) (2.5% w/v) with 0.2% (w/v) of LAP (lithium phenyl-2,4,6-trimethyl-benzoylphosphinate). Thirty microliters of the acini/GelMA suspension were spread on a gold sputtered donor glass slide (30 mm diameter). The bioink was then printed according to a mesh grid pattern (750 µm distance between each point) on a GelMa 5% w/v with 0.2 % of LAP biopaper and gelled at room temperature. The printed distance between donor glass slide and the biopaper was fixed at 1 mm. Laser pulses, enabling the droplets ejection had the following parameters (1064 nm, 30 ns, 85 µJ), resulting on gold ablations with a diameter of 110 µm. The printed droplets and the biopaper were then polymerized during 30 seconds with a UV led (375 nm, 10 mW/cm2). At last, a 2nd layer of warm GelMa 5% w/v with 0.2 % of LAP (150 µL, 37 °C) was deposited on the surface of the printed sample and the all structure was then polymerized during 2 min (375 nm, 10 mW/cm2). The printed sample was then deposited in 6 well-plates in cell medium and incubated at 37 °C during 4, 7 or 14 days depending on the specified time point. The samples were finally post-fixed with 4% (w/v) PFA for 10 min at room temperature, washed twice with PBS and incubated during 2 hours with 1 µmol/L Lipi-Blue staining (Tebu-bio) [10 time the minimal concentration recommended, necessary considering the 3D GelMA surrounding matrix], washed once again with PBS before being imaged. Confocal imaging of both red and blue channels was performed at 10x magnification for red and blue (SPE-II, Leica), with an average of 18 fields of view per condition. Image analysis were conducted using Imaris software. Quantification of lipid staining volume (blue) and total acini volume (red) were extracted exploiting the software's surface reconstruction algorithm. The final quantification was then obtained for each field of view by the normalization of the measured volume of lipids to the measured volume of acini.

Isolation of murine pancreatic cells for single cell RNA sequencing

Pancreas from mice were collected and incubated in multi tissue dissociation kit 1 (Miltenyi Biotec) enzyme cocktail in gentleMACS$^{TM}$ C tubes. Organs were digested with the gentleMACS$^{TM}$ Octo Dissociator (Miltenyi Biotec) during 20 minutes at 37°C following the Cus_37C_m_Panc program. The cell suspension was filtered through a 40 µm strainer and washed with DMEM containing 10% of FBS. Debris were removed using the debris removal solution (Miltenyi Biotec) following the manufacturer's protocol for small volumes. Cells were resuspended in order to obtain a concentration of 1000 cells per µL.

scRNASeq

Murine pancreatic single cells were portioned individually and barcoded using the Chromium Controller and the Chromium Next GEM single cell 3' v3.1 kit (10x Genomics, Inc.). For each condition (CTL or V34), a total of 40000 cells obtained from 7 mice were loaded in different batches. Dual-indexed single-cell RNAseq libraries were prepared according to the manufacturer's instructions. The libraries were profiled with the HS NGS kit for the Fragment Analyzer (Agilent Technologies) and quantified using the KAPA library quantification kit (Roche Diagnostics). The libraries were sequenced on the Illumina NextSeq 550 instrument using High Output 150 cycles kits v2 with following sequencing parameters 28, 10, 10, 90 cycles (read 1, index i7, index i5, read 2). Data will be available on GEO database upon article release.

scRNA-seq data analysis

An average depth of ~75,000 reads/cell was obtained. After sequencing, the output files (bcl2) were converted to FASTQ using cellranger-mkfastq™ software (10x Genomics, Pleasanton, CA, USA). Then to build the (cell, gene) expression matrix we used the cellranger-count algorithm and the GRCh38 human transcriptome reference. Preprocessing, quality control and normalization of Unique Molecular Identifier (UMI), and quality control were assessed using the R package Seurat (*6*, *7*). Cell types were validated using PanGlao https://panglaodb.se/index.html and pancreas_CellMarkerDB https://hpap.pmacs.upenn.edu/ databases, next validated by selective genes (acinar: *Cela3b*, Endothelial: *Cdh5*, Mesenchymal: *Col1a1*, B-cells: *Cd79a*, Erythrocyte: *Hba-a2*, NK: *Nkg7*, Duct: *Onecut1*, Macrophage/Monocyte: *Lyz2*, T cell: *Cd3g*). The single-cell signature scores were scored using the Single-cell Signature-Scorer (*8*). Unless specified otherwise and listed in Table S3, the gene signatures were from MSigDB (https://www.gsea-msigdb.org/gsea/ msigdb/collections.jsp). All scores matrixes were then merged with their correspondents' cell UMAP coordinates, and visualized using Single-Cell Signature-Explorer (*8*) or Multilayer-Viewer (*9*) (open-source software developed at CRCT, Toulouse, France).

Isolation of murine pancreatic cells for single cell RNA sequencing: comparison between the classical protocol and the Miltenyi Biotec protocol (corresponding to the supplementary figure S6)

Pancreas from mice collected and digested using two different protocols. For the classical digestion, the pancreas is cut into 1-3 mm pieces then incubated at 37°C during 30 minutes in digestion solution containing collagenase II 200 U/mL (Sigma-Aldrich), HEPES 0.01M (15630056, Invitrogen), trypsin inhibitor 0.25mg/mL (Sigma-aldrich), and HBSS with $Ca^{2+}/Mg^{2+}$ (H8264, Sigma-Aldrich). Every 5 minutes, the pancreas is manually digested by pipetting using decreasing volumes pipettes. Cells were centrifuged and washed twice with a washing solution containing HBSS without $Ca^{2+}/Mg^{2+}$ (H6648, Sigma-Aldrich), 5% FBS, HEPES 0.01M (15630056, Invitrogen) and trypsin inhibitor 0.1mg/mL (Sigma-Aldrich). Cells are passed through a 100 µm strainer. For the Miltenyi Biotec digestion, the pancreas is incubated in multi tissue dissociation kit 1 (Miltenyi Biotec) enzyme cocktail in gentleMACS™ C tubes. Organs are digested with the gentleMACS™ Octo Dissociator (Miltenyi Biotec) during 20 minutes at 37°C following the Cus_37C_m_Panc program. The cell suspension is filtered through a 40 µm strainer and washed with DMEM containing 10% of FBS.

Cells are analysed by flow cytometry and cell viability was assessed with a Viobility™ 405/452 fixable dye (130-130-403, Miltenyi Biotec).

AR42J transdifferentiation

To induce transdifferentiation, AR42J were treated with DMEM/F12 containing 1µM of Dexamethasone (#D1756, Sigma-Aldrich) for 3 days and the medium was then replaced and supplemented with 20 ng/ml of EGF (236-EG-200, R&D systems) for 5 days. When cells were treated with 0.01 or 0.1 of VPS34 inhibitor SAR405, treatment was applied at the same time than the transdifferentiation treatment and DMSO 0.001% was used as a control.

Western Blot in AR42J or tissue samples

For WB, AR42J cells were treated 2h with VPS34-specific inhibitor SAR405 or Vps34-IN1 at 1, 5 or 10 µM. 0.1% DMSO was used as a control. Isolated acinar cells (see "acini isolation for primary culture" section) were treated with chloroquine at 10 or 25 µM or spautin 2.5 µM at or UCL-TRO-1938 at 10µM or BYL-719 (alpelisib) at 1 µM or transduced with Lv-VPS34. Time points for mouse sample or primary cells are described in the figure legends. For WB on pancreas, a part of snap freeze organ was cut then smashed with pylon and mortar. Proteins were extracted using RIPA lysis buffer (50mM Tris at pH 7.4, 150 mM NaCl, 1% Triton, 1 mM EDTA, 2 mM DTT, 2 mM NaF, 4 mM Na orthovanadate and a protease inhibitor cocktail provided by Roche). 25 µg of proteins were separated by SDS-PAGE and transferred to a nitrocellulose membrane (Amersham). Proteins were detected using primary and secondary antibodies according to table below and were visualized using ChemiDoc (BioRad).

Table related to antibodies used in IF

| Primary antibody | Origin | Dilution | Provider | Secondary antibody |
|---|---|---|---|---|
| LC3B | rabbit | 1/600 | Cell Signaling | 1/1000 |
| LAMP-1 LY1C6 | mouse | 1/100 | Invitrogen | 1/1000 |

Table related to antibodies used in IHC

| Primary antibody | Origin | Dilution | Provider | Secondary antibody |
|---|---|---|---|---|
| VPS34 | Rabbit | 1/200 | Invitrogen/ Thermo Scientifc | Immpress anti rabbit |
| Ki67 | rabbit | 1/1000 | abcam | Immpress anti rabbit |
| REG1 | sheep | 1/500 | R&D systems | 1/200 |
| REG2 | rat | 1/500 | R&D systems | 1/50 |
| REG3A | mouse | | R&D systems | Immpress anti mouse |
| CK19 | rabbit | 1/1000 | Epitomics #3863 | Signal Boost |
| α-amylase | rabbit | 1/3000 | Sigma A8273 | Signal Boost |

Table related to antibodies used in WB

| Primary antibody | Origin | Dilution | Provider | Size (kDa) | Secondary antibody |
|---|---|---|---|---|---|
| p62/ SQSTM1 | rabbit | 1/1000 | Cell Signaling (23214) | 62 | 1/4000-1/5000 |
| LC3B | rabbit | 1/1000 | Cell Signaling (2775S) | 14-16 | 1/10000 |

| | | | | | |
|---|---|---|---|---|---|
| EEA1 | rabbit | 1/1000 | Cell Signaling (3288) | 170 | 1/5000 |
| APPL1 | rabbit | 1/1000 | Cell Signaling (3858) | 82 | 1/5000 |
| Rab5 | rabbit | 1/1000 | Cell Signaling (3547) | 25 | 1/10000 |
| VPS34 | rabbit | 1/500-1/1000 | Cell Signaling (3811) | 100 | 1/10000 (tissue) 1/1000(primary cells) |
| VPS15 | rabbit | 1/500-1/1000 | Abcam (ab128903) | 150 | 1/10000 (tissue) 1/1000(primary cells) |
| Beclin-1 | rabbit | 1/1000 | Cell Signaling (3738) | 52 | 1/10000 |
| Atg14 | rabbit | 1/1000 | MBL (PD026) | 65 | 1/10000 (tissue) 1/2000(primary cells) |
| OPA1 | rabbit | 1/1000 | Cell Signaling (80471) | 80-100 | 1/5000 |
| pDrp1 S616 | rabbit | 1/1000 | Cell Signaling (3455) | 78-82 | 1/5000 |
| pDrp1 S637 | rabbit | 1/1000 | Cell Signaling (4867) | 78-82 | 1/5000 |
| MFN2 | rabbit | 1/1000 | Cell Signaling (9482) | 80 | 1/5000 |
| REG1 | sheep | 1/2000 | R&D systems (AF1657) | 14 | 1/10000 |
| REG2 | rat | 1/500 | R&D systems (MAB2098) | 14 | 1/10000 |
| REG3A | mouse | 1/500 | R&D systems (MAB1745) | 14 | 1/10000 |
| pAkt S473 | rabbit | 1/1000 | Cell Signaling (4060) | 60 | 1/1000 |
| Total Akt | rabbit | 1/1000 | Cell Signaling (4691) | 60 | 1/10000 |
| PARK7 | rabbit | 1/1000 | Abcam (AB18257) | 24 | 1/5000 |
| β-Actin | mouse | 1/1000 | Sigma-Aldrich (AC74) | 42 | 1/10000 |

Tables: List of antibodies for Immuno-fluorescence, Immuno-Histology and Western Blot

RNA extraction and RT-qPCR

Acini were collected by classical digestion as described in "Isolation of murine pancreatic cells for single cell RNA sequencing" section. The cell pellet was resuspended in Trizol (Invitrogen) and RNA was extracted with Direct-zol RNA MiniPrep Plus kit according to manufacturer's protocol (ZymoResearch). 1 µg of RNA was retrotranscribed using iScript cDNA synthesis Kit (BioRad). Primers were designed with Primer-BLAST (NCBI) or taken from the litterature and are listed in Table S1. qPCRs were performed using the SsoFast EvaGreen supermix (Bio-Rad) using StepOne Plus instrument (Thermofisher Scientific). Expression was normalised with Actin, 18S or Rplp0 and represented as fold change using the 2-ΔΔCT formula.

| Genes | Forward primer (5'-3') | Reverse primer (5'-3') | Species | Source |
|---|---|---|---|---|
| **Lipid metabolism genes** | | | | |
| **Acly** | AAAGCTTGGCCTCGTCGG | GGGACGAAGGGTTCAATGAGA | Mouse | Supp. Data Ducheix, PloS One, 2017 PMID 28732092(*10*) |
| **Acaca** | TTACAGGATGGTTTGGCCTTTC | CAAATTCTGCTGGAGAAGCCAC | Mouse | |
| **Acacb** | CCTGAATCTCACGCGCCTA | CAGATGGAGTCCAGACATGCTG | Mouse | |
| **Fasn** | AGTCAGCTATGAAGCAATTGTGGA | CACCCAGACGCCAGTGTTC | Mouse | |
| **Elovl6** | TCTGATGAACAAGCGAGCCA | TGGTCATCAGAATGTACAGCATGT | Mouse | |
| **Scd1** | AGCCTGTTCGTTAGCACCTT | TATCCATAGAGATGCGCGGC | Mouse | Primer Blast |
| **Mitochondrial genes** | | | | |
| **Polg** | CCTGCTCTGGAAGAAGGTGG | CATTCTCGGAGGAGGGCATT | Mouse | Supp. Data Iershov, Nature Comm, 2019, PMID 30952952(*11*) and Primer Blast |
| **Polrmt** | GCTGCCTACATTTCCCACCT | GTGCGGCGTAATGCTGTAAG | Mouse | |
| **Tfb2m** | GCATGTAAGAAAGCGGCCAG | CCACTCTGGCACCAGCTTTA | Mouse | |
| **ATP6** | CCTTCCACAAGGAACTCCAATTTCAC | CTAGAGTAGCTCCTCCGATTAGGTG | Mouse | |
| **ATP8** | ATGCCACAACTAGATACATCAACATG | GTGATTTTGGTGAAGGTGCCAGTGG | Mouse | |
| **ND4** | CACAACACACACCTTAGACGCTTC | CAGCAATTGGAGCTTCAACATGGG | Mouse | |
| **MTCO1** | GCAGGAGCATCAGTAGACCTAAC | GGAGTTTGATACTGTGTTATGGCTGG | Mouse | |
| **ND1** | CACCTACCCTATCACTCACACTAGC | GGCTCATCCTGATCATAGAATGGAG | Mouse | |
| **CYTB** | CTACTGTTCGCAGTCATAGCCAC | CCAATATATGGGATGGCTGATAGGAG | Mouse | |
| **MTCO3** | GGCCACCACACTCCTATTGT | AGGTCAGCAGCCTCCTAGAT | Mouse | |
| **ND3** | GTTGCATTCTGACTCCCCCA | GGTAGACGTGCAGAGCTTGT | Mouse | Primer Blast |
| **MTCO2** | ACCTGGTGAACTACGACTGC | GGACTGCTCATGAGTGGAGG | Mouse | |
| **Autophagy genes** | | | | |
| **BECN1 (beclin1, ATG6)** | GAAACCAGGAGAGACCCAGG | GACATCATCCTGGCTGGGG | Mouse | own design by Primer blast |
| **p62** | AGGAAGCTGCCCTATACCCA | AACCCATGGACAGCATCTGG | Mouse | |
| **ATG5** | TGCATCAAGTTCAGCTCTTCCT | CGCATCCTTGGATGGACAGT | Mouse | |
| **ATG7** | TTGTAGCACCTGCTGACCTG | CCTGGAGCCACCACATCATT | Mouse | |
| **MAP1LC3B** | AAGAGTGGAAGATGTCCGGC | TCGCTCTATAATCACTGGGATCT | Mouse | |
| **MAP1LC3A** | CACACCCATCGCTGACATCT | TGGGAGGCGTAGACCATGTA | Mouse | |
| **GABARAP** | CTGAGGGCGAGAAAATCCGA | CGAGCTTTGGGGGCTTTTTC | Mouse | |
| **GABARAPL1** | CAACAACACCATCCCTCCCA | CTTCCTCGTGGTTGTCCTCA | Mouse | |
| **GABARAPL2** | TCTGGAACACAGATGCGTGG | TTCCACGATCACCGGAACTC | Mouse | |
| **ULK1** | CTCCCCAAGTGGGAACCATC | CATCAGCTCCTTGTGGGGAG | Mouse | |
| **ULK2** | ACGTGCCTTATGGTGCTTCA | AAGGTGTCTGTGTGCTCTCG | Mouse | |
| **ATG4A** | ATCACACTGGCCTCCCTTTG | AATTCTGCCCCTGTGGTTGT | Mouse | |
| **ATG4B** | GCTTCGGTGTGGACAGATGA | TCCACCTCCAATCTCGACCT | Mouse | |
| **ATG4C** | TGTTGCCTGCAAGATCAGGA | TCTTCCCTGTAGGTCAGCCA | Mouse | |
| **VPS34** | CGGCTCAGCAGACCTTTGTA | TGCGGTTCCCACTTTCTCTC | Mouse | |
| **UVRAG** | AGCTGAGATACCAGCATGGC | TGTCTCTTCGGGACAGGGAT | Mouse | |
| **Ambra1** | CCCAGGACTTCCGCTTACAT | GTGTGGCAAGAGGGAAGGAA | Mouse | |

| | | | | |
|---|---|---|---|---|
| **ATG14L (Barkor)** | GCAGGTCAGGACCCTTTGAA | AGGTCAGTCTCCTCATCGCT | Mouse | |
| **WIPI1 (ATG18)** | CACAGGACGCCTGAACTTCT | GGCAGTTTCTGGATCGTGGA | Mouse | |
| **ATG2A** | TTGGATGTGCTGGATGACCC | GTAGAGGGGCCTGTAGTCCA | Mouse | |
| **ATG3** | GTGGCAGCTGGAGATCACTT | ACACCGCTTGTAGCATGGAA | Mouse | |
| **ATG9** | GCTATCCCTGTGCTACACCC | GATCTGTGATGCCCCCGTAG | Mouse | |
| **ATG10** | CGATGGCTGGGAATGGAGAA | GGCATGTGGTGTCAAGGTCT | Mouse | |
| **ATG12** | TAAACTGGTGGCCTCGGAAC | ATCCCCATGCCTGGGATTTG | Mouse | |
| **FIP200 (Rb1cc1)** | CACAGTCAGCTGCTTCTCCA | CACTGCCGGACATAAGGTGT | Mouse | |
| **DRAM1** | CTCAGGGGAATGGCTTTCGT | GAGATGATGAAGGCTGCGGA | Mouse | |
| **CTSD (cathepsin D)** | TTGTGAGAAGGTGTCCAGCC | CATGAAGCCACTCAGGCAGA | Mouse | |
| **Bcl2** | CAGCATGCGACCTCTGTTTG | TCACTTGTGGCCCAGGTATG | Mouse | |
| **STK38** | ACGCTCGGAAGGAAACAGAG | CATTGCGTACACATGCCCTG | Mouse | |
| **RalB** | GGAGAAGGCTTTCTGCTGGT | GCAGTGGGATCTTGTCCTCC | Mouse | |
| **RalA** | AAGCGGACAGCTACAGGAAG | ATCTGCACTTCTTCCCCGTC | Mouse | |
| **Acinar and ductal genes** | | | | |
| **Ck19** | ACACCAGGCATTGACCTAGC | ACCAGGCTTCAGCATCCTTC | Rat | Primer Blast |
| **Ck7** | CCGGAATGAGATTGCGGAGA | ACGGTGTCAATCTCTGCCTG | Rat | |
| **Mist1** | CCTCACACTGGCCAAGAACT | GCGGCTGCTGGACATAGTAA | Rat | |
| **Ptf1a** | CTACGAAAAGCGCCTCTCCA | TAGCCTATGGCTAAGCGCAG | Rat | |
| **Amy2a** | TCTCCCAAGGAAGCAGACCT | ACGCGGCCATTTCCAAAGTA | Rat | |
| **Ctrc** | TGGGGGAAGTCTCATCACCA | TATACTTCCCCAGGCCCACA | Rat | |
| Homeostasis genes | | | | |
| **Gapdh** | GACGGGCCATCAAAGAAGGA | GCTCTGGGATGACTTTGCCT | Rat | Primer Blast |
| **Rplp0** | CGAAGCCACACTGCTGAACA | ACCCTCTAGGAAGCGAGTGT | Rat | |

Table: List of primers.

Live and dead assay

Eighty thousand AR42J cells were plated in 6-well plates and transdifferentiated as previously described. Cells were treated with 0.01 or 0.1 of VPS34 inhibitor SAR405 at the same time than the transdifferentiation treatment and DMSO 0.001% was used as a control. Living and dead cells were labeled for 1 h using the LIVE/DEAD® kit (ThermoFisher Scientific) containing 2 µM calcein and 4 µM EthD-1 (Ethidium homodimer-1). Cells were observed and imaged with an AxioVert microscope after 8 days using ZEN® software (Carl Zeiss). The percentage of living or dead cells was counted and represented as the percentage of the whole population.

BODIPY staining

Eighty thousand AR42J cells were plated in 6-well plates with sterile coverslips. Cells were treated with 0.01 or 0.1 of VPS34 inhibitor SAR405 and DMSO 0.001% was used as a control. After 4 or 8 days of culture, cells were fixed 10 minutes with paraformaldehyde (PFA) 4% in PBS. They then were washed with PBS and incubated with 70µl/well of a solution containing BODIPY 493/503 (1 mg/mL, Invitrogen), beforehand diluted at 1/1000 in 150mM NaCl, in the dark at room temperature for 10 minutes. Cells were then washed and covered with 1 mL of a solution of DAPI (1 mg/ml, Thermofisher) previously diluted at 1/10000 in PBS, for 1 minute. Cells were mounted with Mowiol (Sigma-Aldrich) on a glass slide and then observed with a Cell Observer video microscope (Zeiss) with a 63x objective. 10 pictures were taken per slide and the number of vacuoles was counted using ImageJ. For positive control conditions, Oleate (1/100, Sigma-Aldrich) was added to the medium 24 hours before the BODIPY staining.

Acini isolation for primary culture

Pancreas were collected from mice and washed by cold HBSS 1X without Ca2+ and Mg2+ solution (H6648, Sigma-Aldrich /Merk) containing collagenase II (C6885- Sigma-Aldrich /Merk). Pancreases were immediately mechano-chemically digested by several ups and downs with decreasing volume of pipettes in HBSS 1X with Ca2+ and Mg2+ solution (H8264, Sigma-Aldrich /Merk) containing collagenase II (C6885- Sigma-Aldrich /Merk) and trypsin inhibitor at 37°C for 30 minutes. Following several washes by HBSS 1X without Ca2+ and Mg2+ solution containing HEPES (15630056, Invitrogen ThermoFisher scientific), trypsin inhibitor and FBS (CVFSVF00-01, Eurobio) each followed by centrifugation for 1 minute at 400 rpm 4°C, acini were collected using 100µM strainer into DMEM, high glucose, GlutaMAX(TM) (61965, Gibco, Invitrogen ThermoFisher scientific) and next used in culture.

Immunofluorescence for isolated acini

Acinar cells within 8-ibidi plates were fixed/permeabilized in a solution of methanol 50% (v/v) and acetone 50% (v/v) previously cooled for 10 min at -20°C. Following washing, cells were blocked with 3% BSA/PBS for 1 h at RT. Then, wells were washed with PBS and incubated overnight at 4°C with the primary antibody diluted in solution 3% BSA/PBS. Wells were then washed in PBS and incubated with secondary antibody diluted in 3% BSA/PBS for 1h at RT. Wells were washed with PBS and stained with Hoechst 33342 (ab228551, abcam) or DAPI (D9542, Sigma-Aldrich/Merk) diluted in fluoromount-G (00-4958-02, ThermoFisher). Cells were imaged under a confocal microscope using ZEN blue software. Quantitative analysis was performed using ImageJ software.

Immunohistochemistry

Formalin-fixed (Sigma HT501128), paraffin-embedded pancreas sections (4 µm) were dewaxed by incubation at 60°C for 1 hour followed by xylene and rehydrated through graded alcohols. Pircrosirius red stainings (ab150681) were performed according to (*12*). Slides were analysed under light and polarized light microscopy (AxioObserver ZI ZEISS). Tissue sections were pressure cooked in citrate buffer for 12 min, incubated with PBS, 3% H2O2 (H1009, Sigma-Aldrich) 10% methanol (32213, Sigma-Aldrich) for 15 min at RT to block endogenous peroxidase and then non specific binding sites were blocked by Protein Block (X0909, DAKO) for 1 hr at RT within humid chamber. Slides were incubated with the primary antibodies diluted in antibody diluent (S2022, DAKO) overnight at 4°C within humid chamber. Then, slides were incubated with secondary antibodies linked with HRP or biotinylated antibody (Vectastain, PK-7200, Vector Labs) or using the HRP-Detection Reagent SignalStain® Boost (CST 8114), diluted in antibody diluent, for 1h at RT. Secondary antibodies were revealed using DAB (SK-4105, Vector Labs) or AEC Substrate Chromogen (Dako K3464) (CK19, amylase). Sections were counterstained with haematoxylin (1.09249, Merck) before mounting on coverslips with glycergel (C0563, DAKO). After scanning with Pannoramic ® 250 Flash III microscope, analyses were done with QuPath Software(*13*). To define areas of interest, common tools of QuPath were used additionally to Segment Anything Model extension(*14*). For REG3A analyse, pixel classifier was used. For Ki67 analysis, positive cell detection was used, more than 10 000 cells were counted for each animal.

Viral transduction

Isolated acinar cells and viral particles were mixed to obtain a Multiplicity of infection (MOI) (refers to the number of viral particle(s) present relative to host cell(s)) of 3 or 5 in 6-wells plates. Polybrene (TR-1003-G, Sigma-Aldrich) was added at a final concentration of 10µg/mL. Plates were shaken each 15 minutes for 1 hour and then each 1 hour for 3 hours.

Statistics

Statistical analyses were performed with GraphPad Prism using student T-test for in vitro data; non-parametric Mann-Whitney for all in vivo data; Mantel-Cox statistical test was used for all survival curves: *p<0.05, **p<0.01, ***p<0.001, ****p<0.0001. Statistical relevance of the murine KC cohort size was determined using a Power Calculation test (www.lasec.cuhk.edu.hk/.../power_calculator_14_may_2014.xls).

Type or paste text here. This should be additional explanatory text, such as: extended technical descriptions of results, full details of mathematical models, extended lists of acknowledgments, etc. It should not be additional discussion, analysis, interpretation, or critique.

| Patient | Pathology |
|---|---|
| Patient#1 | Alcoholic CP |
| Patient#2 | Obstructive CP adjacent to ampullary lesion |
| Patient#3 | Atrophic CP of unknown origin |
| Patient#4 | Obstructive CP adjacent to IPMN (moderate dysplasia) |
| Patient#5 | Obstructive CP adjacent to ampullary PDAC |
| Patient#6 | Obstructive CP adjacent to cephalic PDAC |
| Patient#7 | Obstructive CP adjacent to PDAC |
| Patient#8 | Obstructive CP on PDAC and PanIN3 |

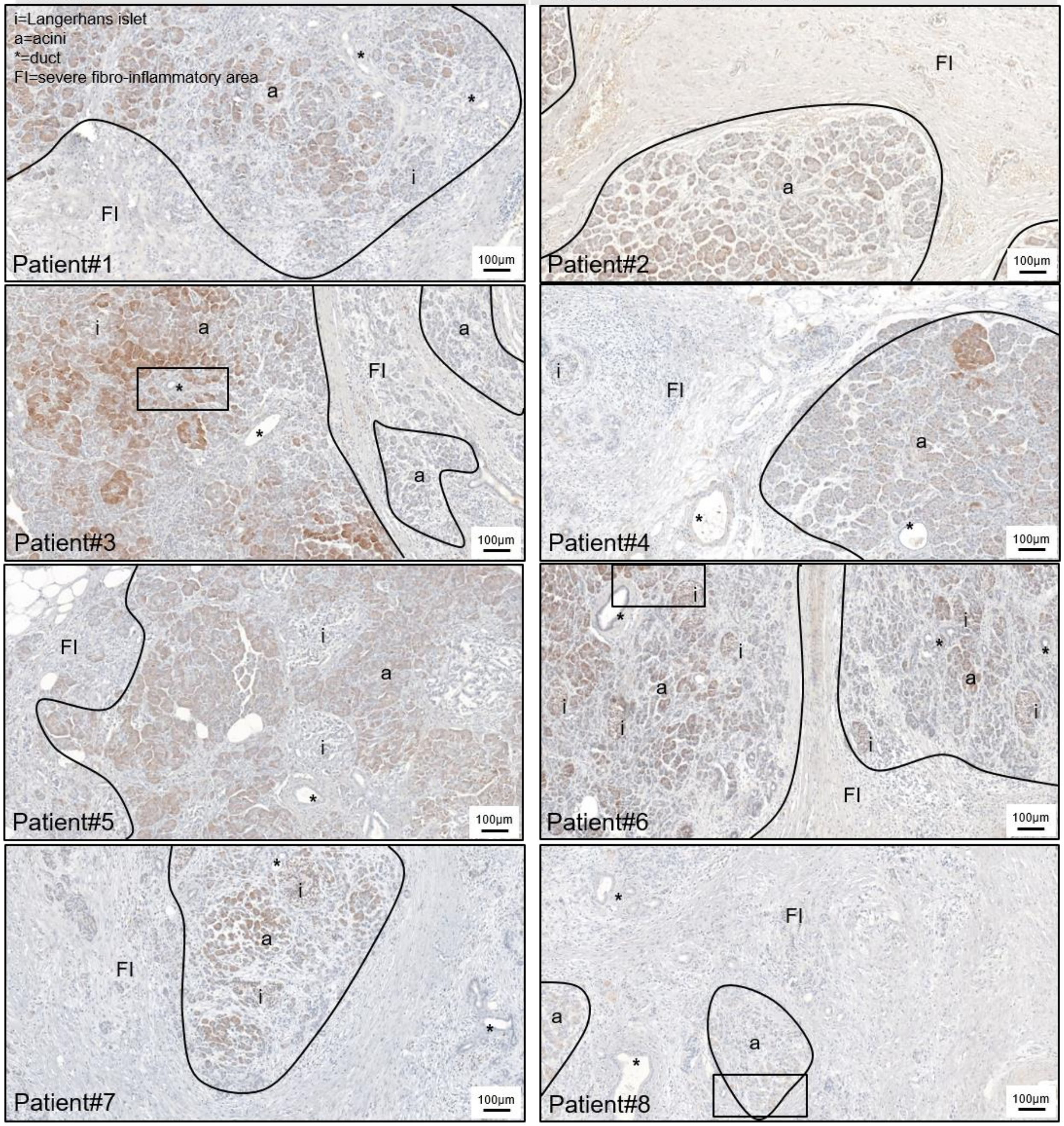


**Fig. S1. Heterogeneity of VPS34 level in healthy and inflamed human pancreas (from**

**chronic pancreatitis patients).**

Pathological context of patients, etiology of chronic pancreatitis (CP) and corresponding immunohistochemistry of VPS34 in human pancreas; a highlights acini areas; i highlights Langerhans islets; asterisk highlights ducts; FI refers to severe fibro-inflammatory area, characterized by fibrosis, loss of acinar tissue and duct changes. N=8. High magnification, insets are shown in Fig. 1.

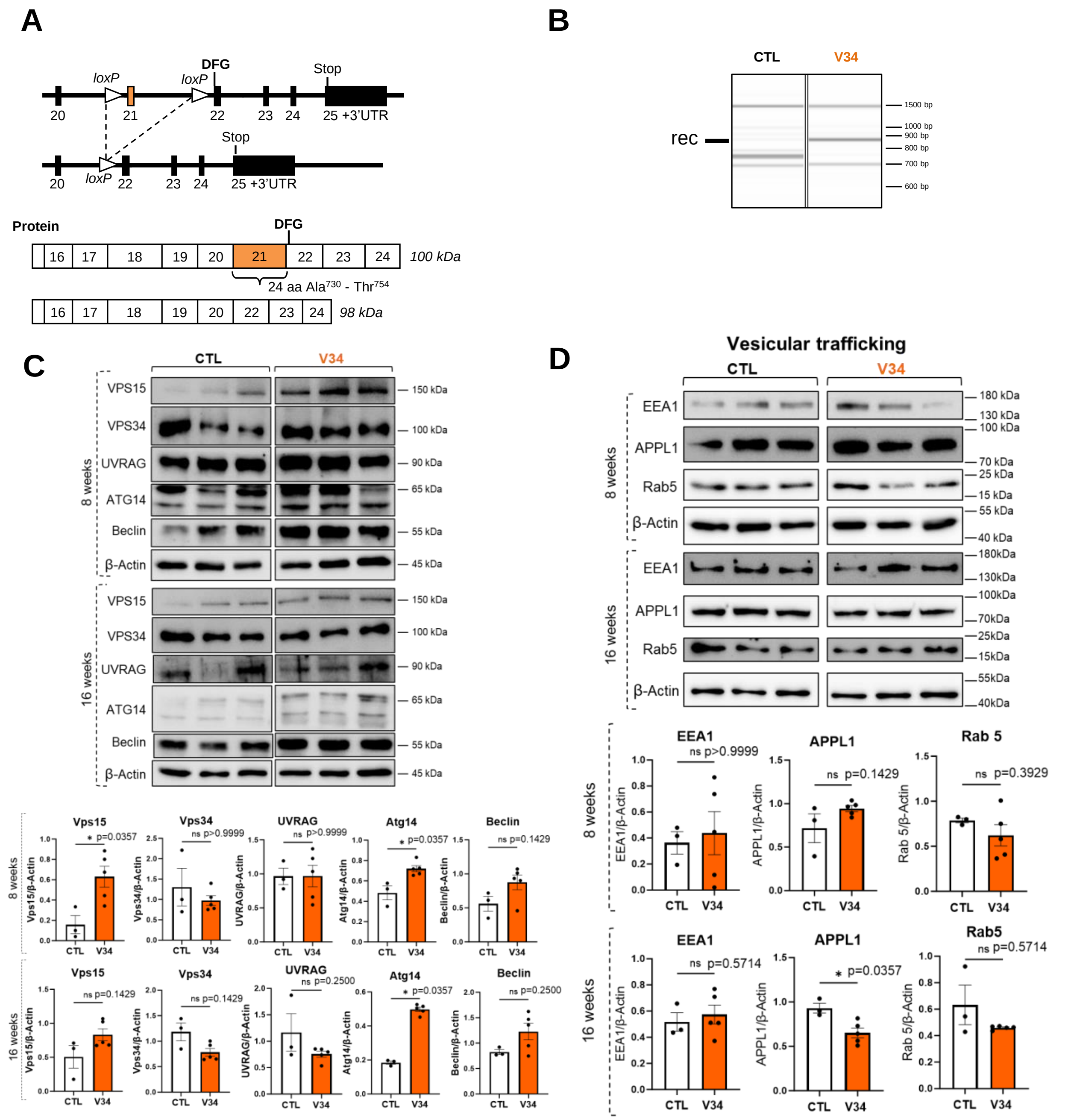


**Fig. S2. Validation of V34 model.**

(**A**) Gene construct of VPS34 floxed mice. In *PIK3C3* gene encoding for VPS34, the kinase domain corresponds to exons 17-26. Deletion of exon 21 will result in the deletion from Ala730 to Thr754 as exons remains in frame for translation. (**B**) PCR of genomic DNA from CTL or V34 2 weeks after tamoxifen injection using non rec and rec primers [Vps34F; Vps34R; Vps34F; Rrec1(R22,5)] primers. A selective band is detected upon recombination only in V34 mice. See supplementary Table 1 for tool details.

**(C)** Western blot analysis showing VPS34 complex I and II members in pancreas tissues from CTL and V34 mice. Protein levels were normalized with β-Actin level. Statistical analysis Mean +/- SEM, (*$p<0.05$, Mann Whitney test), CTL n=3, V34 n=5. **(D)** Western blot analysis showing proteins involved in vesicular trafficking in pancreas tissues from CTL and V34 mice. Protein levels were normalized with β-Actin level. Statistical analysis Mean +/- SEM, (*$p<0.05$, Mann Whitney test), CTL n=3, V34 n=5.

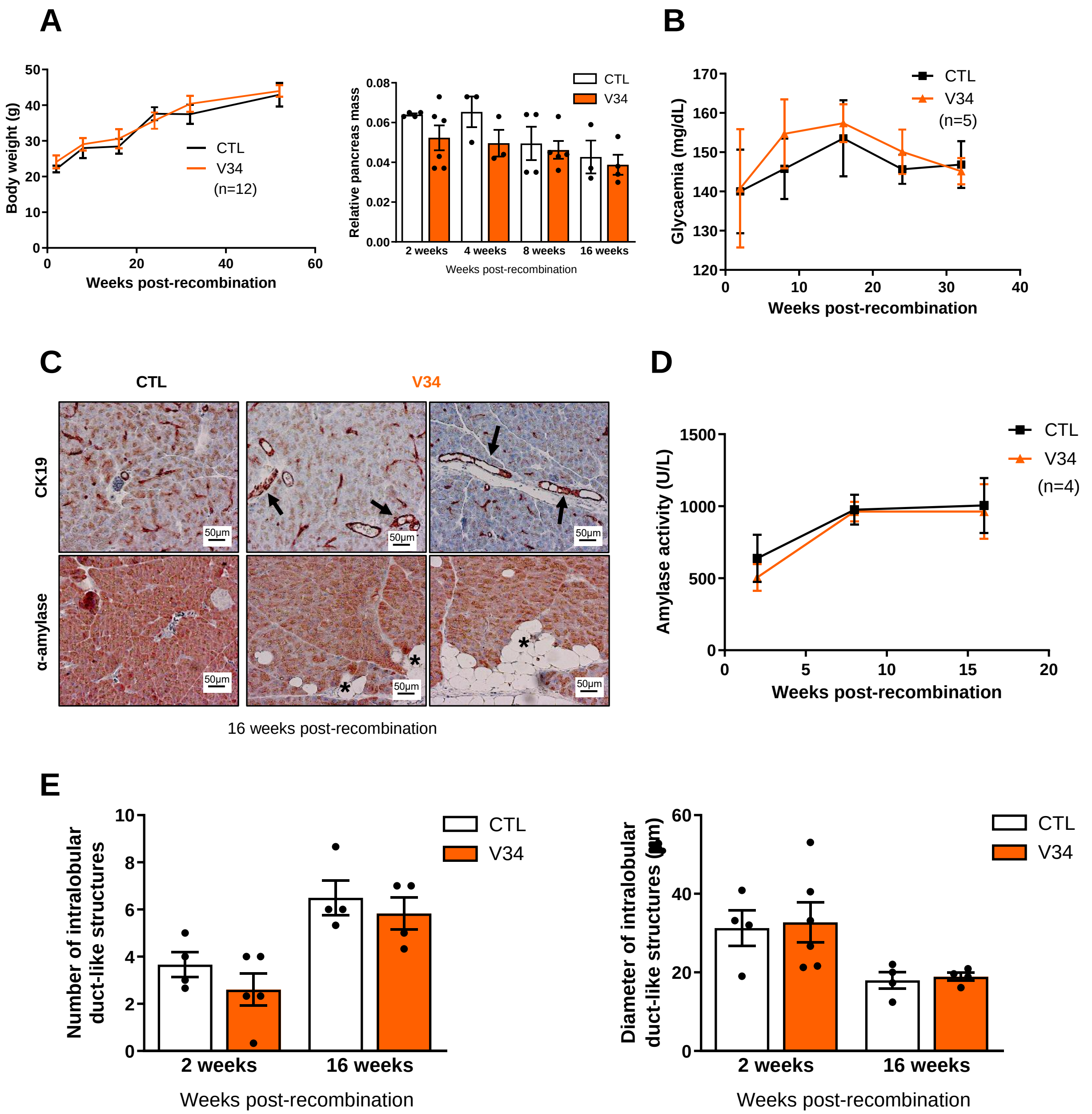


**Fig. S3. General information on V34 mice**

(**A**) Weight evolution and relative pancreas mass of control (CTL) or VPS34lox/lox (V34) mice at different times after recombination. (**B**) Glycaemia (mg/mL) of CTL or V34 mice at different times after recombination. N=5 Mean +/- SEM

(**C**) CK19 and α-amylase immunochemistry on pancreatic sections of CTL or V34 mice 16 weeks post-recombination. (**D**) Quantification of α-amylase staining of CTL or V34 at different times after recombination. N=4 Mean +/- SEM

(**E**) Quantification of the number and diameter (μm) of intra lobular duct-like structures in pancreas sections from CTL or V34 mice 2 or 16 weeks after recombination. N≥4 Mean +/- SEM

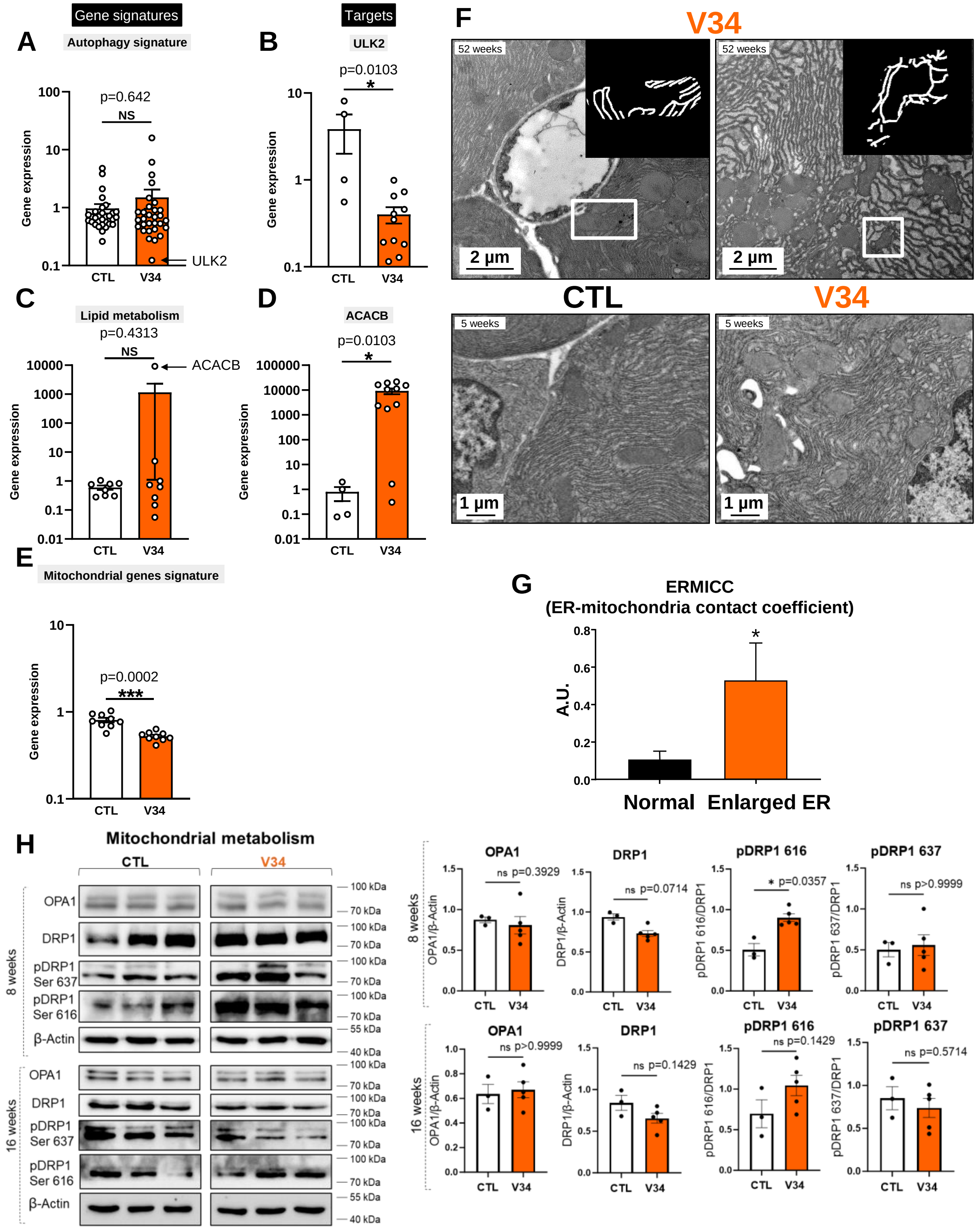


**Fig. S4. Subcellular and molecular characterization of CTL and V34 pancreas.**

**(A-E)** RT-qPCR on isolated acinar cells from CTL (n=4) and V34 (n=11) 52 weeks after tamoxifen injection, analysing the expression of (**A**) macroautophagy signature (30 genes), (**B**)

ULK2, (**C**) lipid metabolism signature (8 genes), (**D**) ACACB and (**E**) mitochondrial gene signature. Mean +/- SEM (*p<0.05, ** p<0.01, *** p<0.001, Student t-test).
(**F**) Electron microscopy pictures of CTL and V34 mice, 5 or 52 weeks after recombination. (**G**) Quantification of endoplasmic reticulum-mitochondria contact coefficient in V34 pancreas at 52 weeks. Mean +/- SEM, (*p<0.05, Student t-test). N=3 independent pancreas
**(H)** Western blot analysis showing proteins involved in mitochondrial metabolism on pancreas tissues from CTL and V34 mice at indicated time point (8 or 16 weeks after tamoxifen). Protein levels were normalized with β-Actin. Mean +/- SEM, (*p<0.05, Mann Whitney test), CTL n=3, V34 n=5.

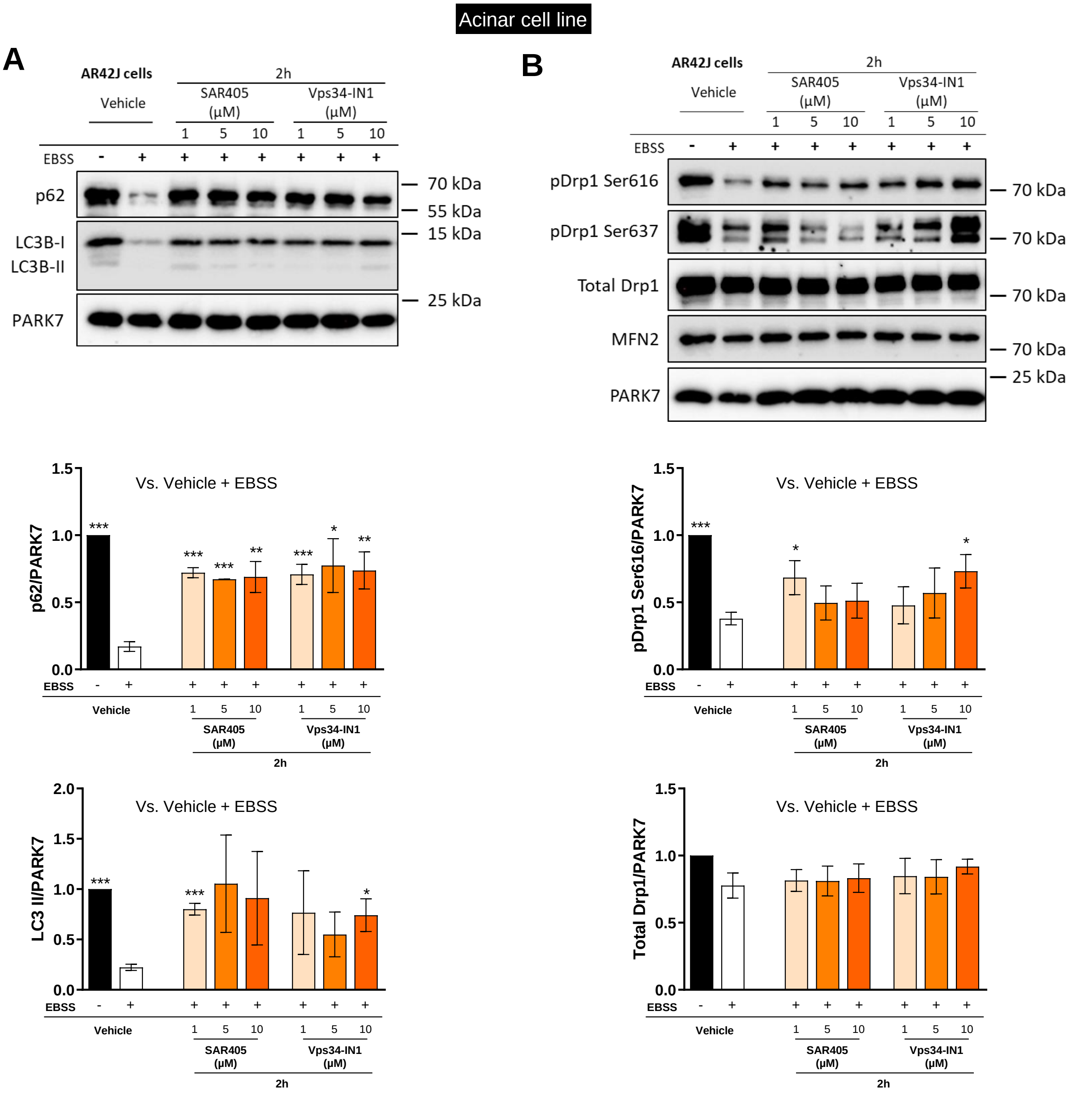


**Fig. S5. VPS34 pharmacological inhibition inhibits autophagy pathway and modulates mitochondrial dynamics.**

(**A, B**) Acinar cell line AR42J was treated with VPS34 pharmacological inhibitor SAR405 or Vps34-IN1 as indicated. Western blot analysis showing proteins involved in (**A**) autophagy/vesicular trafficking and (**B**) mitochondrial dynamics. Mean +/- SEM n=3 (* $p<0.05$, ** $p<0.01$, *** $p<0.001$, Student t-test).

<insert page break here>

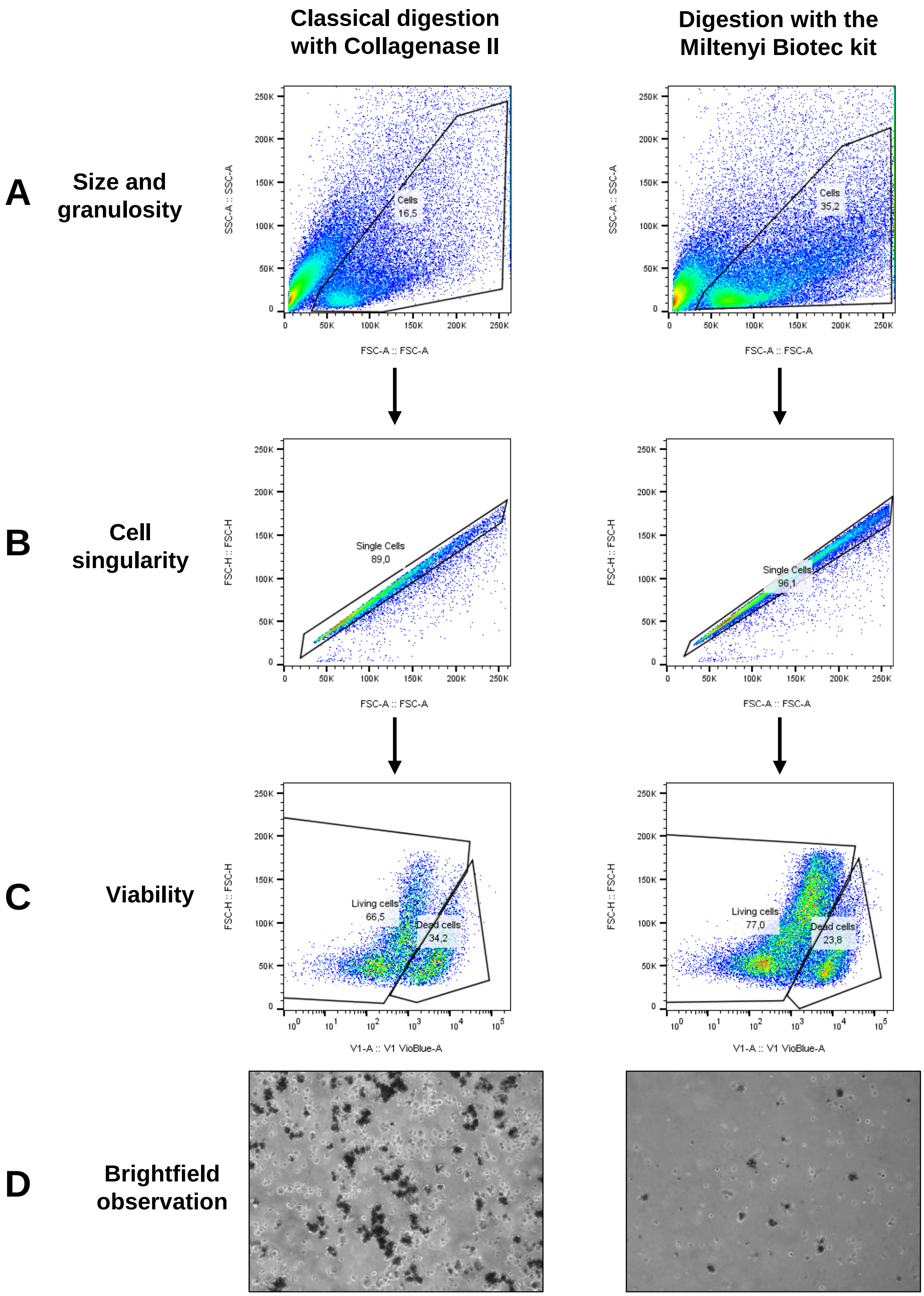


**Fig. S6. Optimisation of mouse pancreas single cell preparation**

Mice pancreas are subjected to classical digestion with collagenase II or with the Miltenyi Biotec kit multi tissue dissociation kit 1. (**A**) Cell preparations are analysed by flow cytometry for size and granularity, (**B**) cell singularity (single cells versus clusters of cells) and (**C**) viability. (**D**) Cell are observed in brightfield.

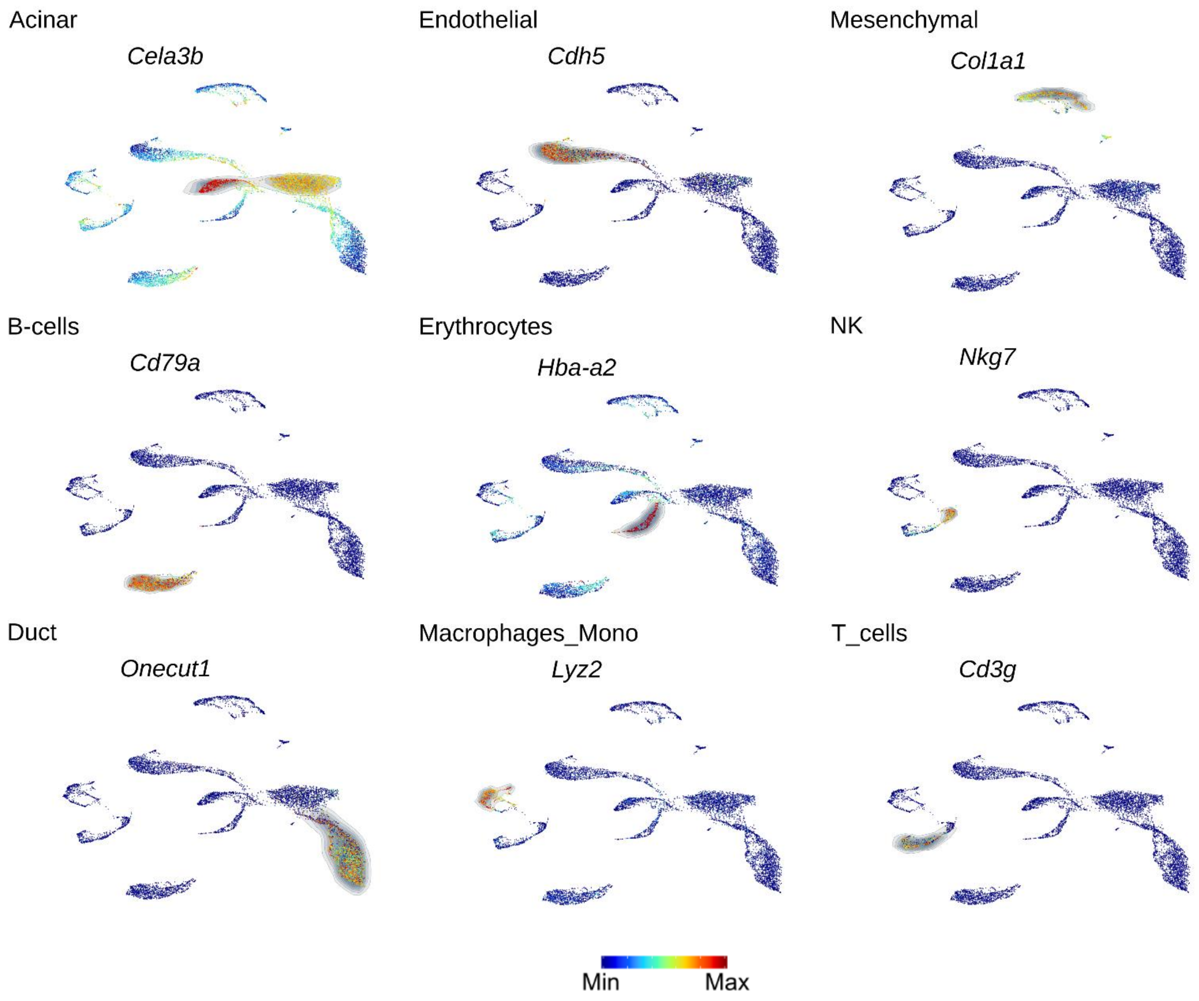


**Fig. S7. UMAP representation of cell type markers.**

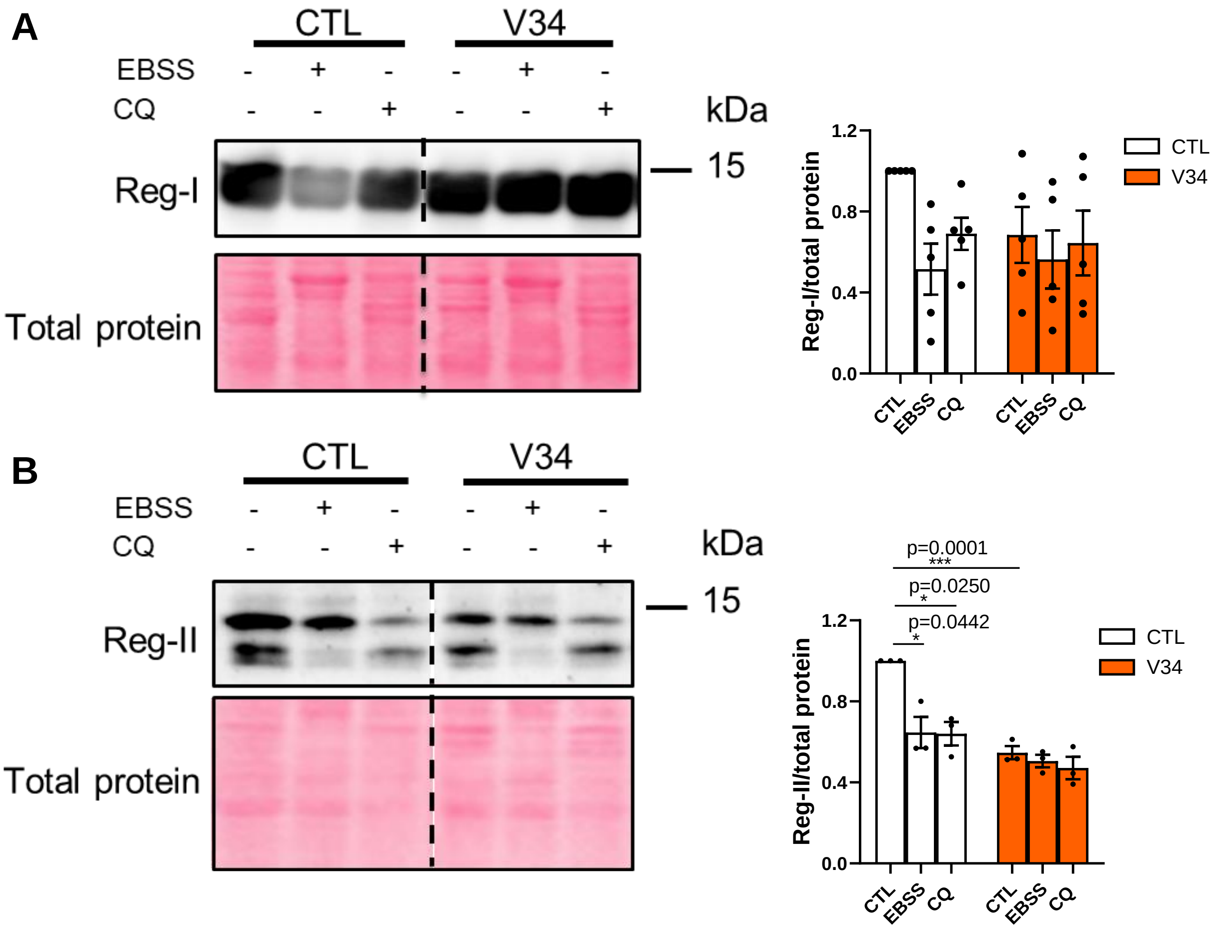


**Fig. S8. Regulation of other REG protein levels in CTL or V34 acinar cells.**

**(A,B)** Western blot analysis showing (A) Reg-I and (B) Reg-II on isolated acinar cells 5-9 weeks after tamoxifen injection from CTL or V34 mice. Isolated primary acinar cells treated with or without EBSS (24 hrs) or chloroquine CQ (10µM; 24 hrs). Protein levels were normalized with total protein. Mean +/- SEM, (*p<0.05, ** p<0.01, *** p<0.001), N=at least 3. Western blot analysis of (A) Reg-I and (B) Reg-II in primary acinar cells treated with or without EBSS or Chloroquine (CQ) from CTL or V34 mice. Protein levels were normalized with total proteins. Mean +/- SEM, (*p<0.05, ** p<0.01, *** p<0.001), N=3.

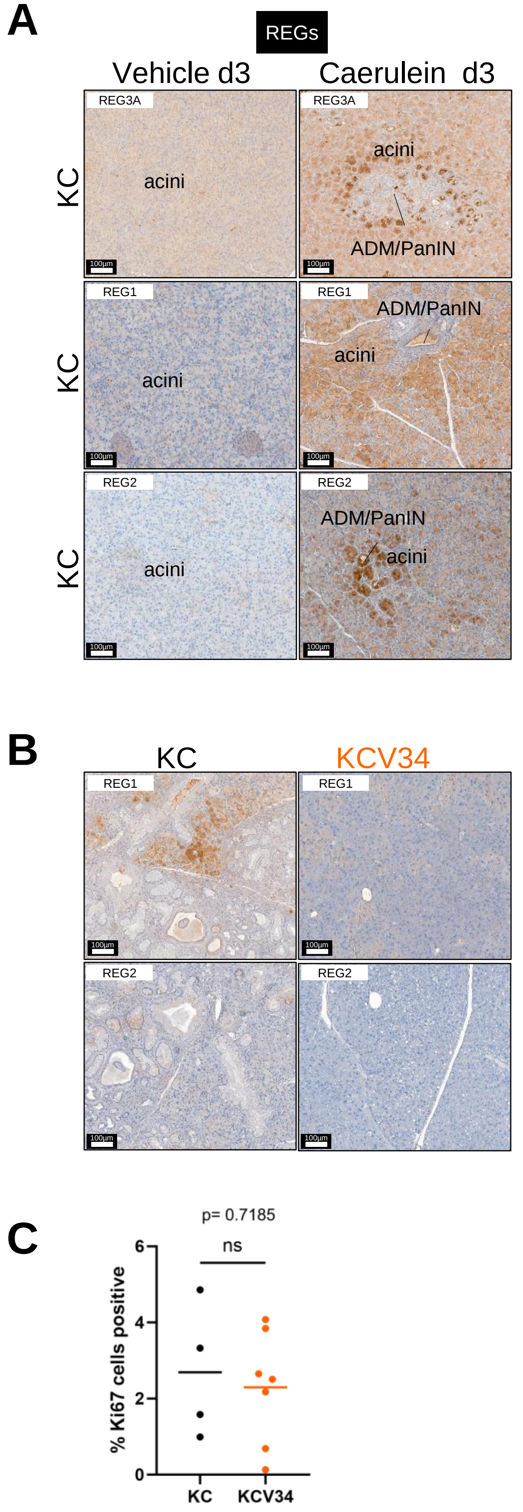


**Fig. S9. Supplementary figure to Figure 4.**

(**A**) Imunohistochemical staining of REG3A, REG1 and REG2 in pancreatic tissues from KC mice 3 days post treatment with vehicle or Caerulein. Scale bar = 100µM.

(**B**) Imunohistochemical staining of REG1 and REG2 in pancreatic tissues from KC and KCV34 mice. Scale bar= 100µM.
(**C**) Imunohistochemical staining of Ki67 in pancreatic tissues from KC and KCV34 mice. Individual values and mean, Student t-test (* $p<0.05$, ** $p<0.01$, *** $p<0.001$); KC n=4; KCV34 n=7.

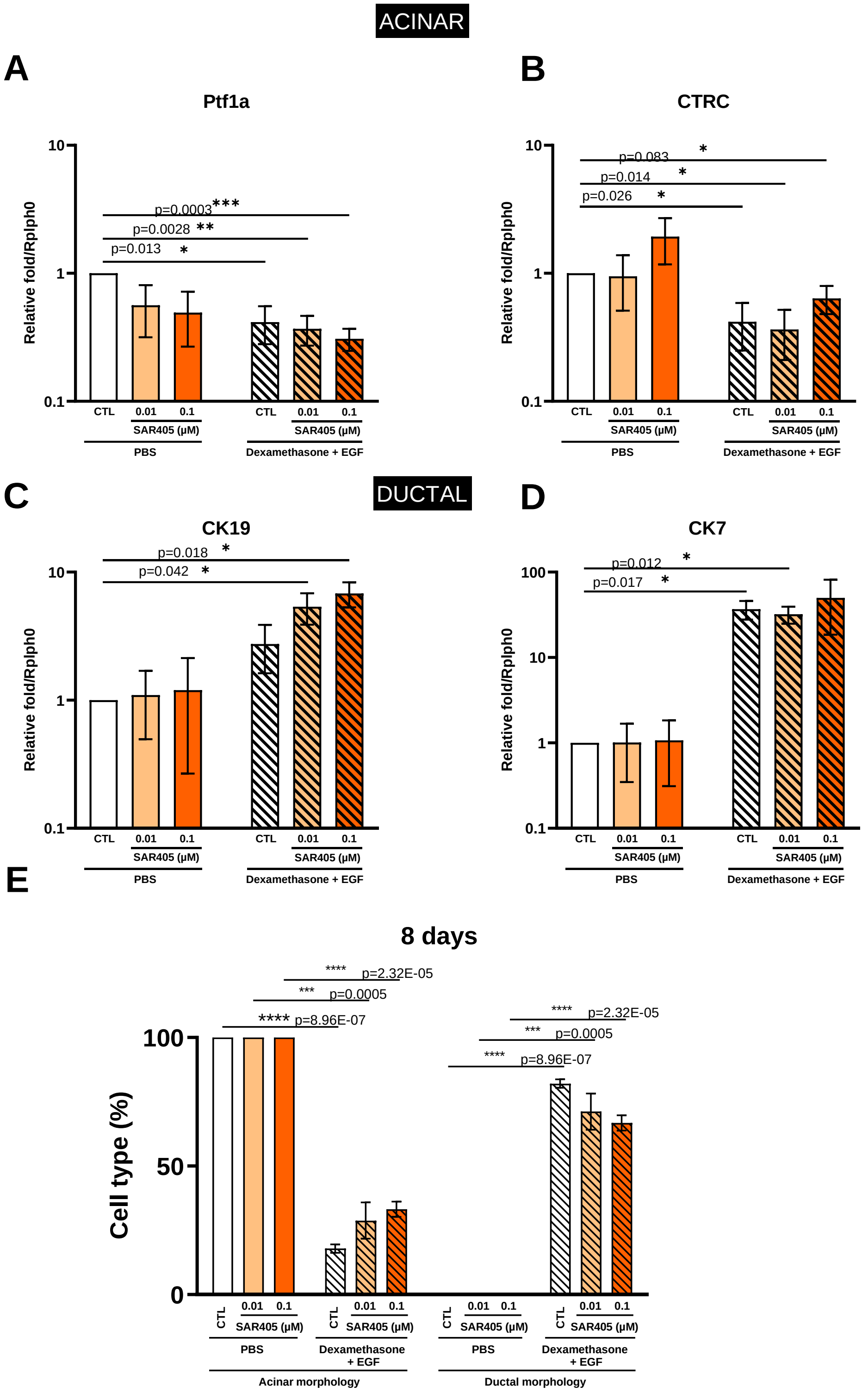


**Fig. S10. VPS34 pharmacological inhibition does not alter acinar-to-ductal transdifferentiation in vitro.**

(**A-E**) Acinar cell line AR42J was treated with VPS34 pharmacological inhibitor SAR405 as indicated and were transdifferentiated or not for 8 days with Dexamethasone and EGF. Relative expression of acinar genes (**A**) Ptf1a and (**B**) CTRC or ductal genes (**C**) CK19 and (**D,** CK7. The expression was normalised with the ribosomal protein Rplp0 expression and represented as fold change using the 2-ΔΔCT formula. Mean +/- SEM (* $p < 0.05$, ** $p < 0.01$, *** $p < 0.001$, N = 3). (**E**) AR42J cells treated as previously described were morphologically observed at the microscope. Acinar or ductal cells were counted and represented as percentages of the whole population. Mean +/- SEM (*** $p<0.001$, **** $p<0.0001$, N = 3).

**Table S1. Detailed List of Material, Tools and protocols**

**Table S2. scRNAseq differential cluster markers with statistical analysis**

**Table S3. List of genes found in specific gene signatures**

### References of Supplementary Materials


1. C. J. F. Courreges, E. C. M. Davenport, B. Bilanges, E. Rebollo-Gomez, J. Hukelmann, P. Schoenfelder, J. R. Edgar, D. Sansom, C. Scudamore, R. Roychudhuri, O. A. Garden, B. Vanhaesebroeck, K. Okkenhaug, Lack of phosphatidylinositol 3-kinase VPS34 in regulatory T cells leads to a fatal lymphoproliferative disorder without affecting their development. [Preprint] (2024). https://doi.org/10.1101/2024.01.08.574346.

2. P.-P. Prévot, A. Simion, A. Grimont, M. Colletti, A. Khalaileh, G. Van den Steen, C. Sempoux, X. Xu, V. Roelants, J. Hald, L. Bertrand, H. Heimberg, S. F. Konieczny, Y. Dor, F. P. Lemaigre, P. Jacquemin, Role of the ductal transcription factors HNF6 and Sox9 in pancreatic acinar-to-ductal metaplasia. *Gut* **61**, 1723–1732 (2012).

3. M. Brunet, C. Vargas, M. Fanjul, D. Varry, N. Hanoun, D. Larrieu, L. Pieruccioni, G. Labrousse, H. Lulka, F. Capilla, A. Ricard, J. Selves, A. Couvelard, V. Gigoux, P. Cordelier, J. Guillermet-Guibert, M. Dufresne, J. Torrisani, The E3 ubiquitin ligase TRIP12 is required for pancreatic acinar cell plasticity and pancreatic carcinogenesis. *J Pathol* **263**, 466–481 (2024).

4. K. N. Diakopoulos, M. Lesina, S. Wörmann, L. Song, M. Aichler, L. Schild, A. Artati, W. Römisch-Margl, T. Wartmann, R. Fischer, Y. Kabiri, H. Zischka, W. Halangk, I. E. Demir, C. Pilsak, A. Walch, C. S. Mantzoros, J. M. Steiner, M. Erkan, R. M. Schmid, H. Witt, J. Adamski, H. Algül, Impaired autophagy induces chronic atrophic pancreatitis in mice via sex- and nutrition-dependent processes. *Gastroenterology* **148**, 626-638.e17 (2015).

5. Q. Chu, T. F. Martinez, S. W. Novak, C. J. Donaldson, D. Tan, J. M. Vaughan, T. Chang, J. K. Diedrich, L. Andrade, A. Kim, T. Zhang, U. Manor, A. Saghatelian, Regulation of the ER stress response by a mitochondrial microprotein. *Nat Commun* **10**, 4883 (2019).

6. T. Stuart, A. Butler, P. Hoffman, C. Hafemeister, E. Papalexi, W. M. Mauck, Y. Hao, M. Stoeckius, P. Smibert, R. Satija, Comprehensive Integration of Single-Cell Data. *Cell* **177**, 1888-1902.e21 (2019).

7. A. Butler, P. Hoffman, P. Smibert, E. Papalexi, R. Satija, Integrating single-cell transcriptomic data across different conditions, technologies, and species. *Nat Biotechnol* **36**, 411–420 (2018).

8. F. Pont, M. Tosolini, J. J. Fournié, Single-Cell Signature Explorer for comprehensive visualization of single cell signatures across scRNA-seq datasets. *Nucleic Acids Res.* **47**, e133 (2019).

9. J.-P. Cerapio, M. Perrier, C.-C. Balança, P. Gravelle, F. Pont, C. Devaud, D.-M. Franchini, V. Féliu, M. Tosolini, C. Valle, F. Lopez, A. Quillet-Mary, L. Ysebaert, A. Martinez, J. P. Delord, M. Ayyoub, C. Laurent, J.-J. Fournie, Phased differentiation of γδ T and T CD8 tumor-infiltrating lymphocytes revealed by single-cell transcriptomics of human cancers. *Oncoimmunology* **10**, 1939518 (2021).

10. S. Ducheix, A. Montagner, A. Polizzi, F. Lasserre, M. Régnier, A. Marmugi, F. Benhamed, J. Bertrand-Michel, L. Mselli-Lakhal, N. Loiseau, P. G. Martin, J.-M. Lobaccaro, L. Ferrier, C. Postic, H. Guillou, Dietary oleic acid regulates hepatic lipogenesis through a liver X receptor-dependent signaling. *PLoS One* **12**, e0181393 (2017).

11. A. Iershov, I. Nemazanyy, C. Alkhoury, M. Girard, E. Barth, N. Cagnard, A. Montagner, D. Chretien, E. I. Rugarli, H. Guillou, M. Pende, G. Panasyuk, The class 3 PI3K coordinates autophagy and mitochondrial lipid catabolism by controlling nuclear receptor PPARα. *Nat Commun* **10**, 1566 (2019).

12. N. Therville, S. Arcucci, A. Vertut, F. Ramos-Delgado, D. F. Da Mota, M. Dufresne, C. Basset, J. Guillermet-Guibert, Experimental pancreatic cancer develops in soft pancreas: novel leads for an individualized diagnosis by ultrafast elasticity imaging. *Theranostics* **9**, 6369–6379 (2019).

13. P. Bankhead, M. B. Loughrey, J. A. Fernández, Y. Dombrowski, D. G. McArt, P. D. Dunne, S. McQuaid, R. T. Gray, L. J. Murray, H. G. Coleman, J. A. James, M. Salto-Tellez, P. W. Hamilton, QuPath: Open source software for digital pathology image analysis. *Sci Rep* **7**, 16878 (2017).

14. K. Sugawara, Training deep learning models for cell image segmentation with sparse annotations. Bioinformatics [Preprint] (2023). https://doi.org/10.1101/2023.06.13.544786.
